# Functional Equivalence of the Proneural Genes *Neurog1* and *Neurog2* in the Developing Dorsal Root Ganglia Highlights the Importance of Timing of Neurogenesis in Biasing Somatosensory Precursor Fates

**DOI:** 10.1101/2025.01.08.631905

**Authors:** Simon Desiderio, Pauline Cabochette, Stephanie Venteo, Gautier Tejedor, Farida Djouad, Patrick Carroll, Fabrice Ango, Alexandre Pattyn

## Abstract

Distinct subfamilies of proneural bHLH transcription factors (TFs) control neurogenesis and influence neuronal fate specification throughout the developing nervous system. While members of different subfamilies generally display divergent expression and functions, those within the same subfamily are often co-expressed, raising questions about their respective roles. Here, we assess the functional equivalence of the two paralogous proneural TFs Neurog1 and Neurog2, focusing on neurons of the dorsal root ganglia (DRG) which underlie our ability to process somatosensory information. During development, distinct classes of DRG neurons -classified as mechano/proprioceptors or thermo/nociceptors-are generated during two successive neurogenic phases, which are respectively controlled by Neurog2 and Neurog1, in line with their dynamic and complementary spatiotemporal expression patterns. Therefore, in this system, each TF appears to play unique roles in regulating neurogenesis and specifying distinct neuronal identities. Yet, using a gene replacement strategy *in vivo*, we show that, despite subtle divergences masked by dose-compensation effects, Neurog1 and Neurog2 are essentially functionally interchangeable in all aspects of DRG development, indicating that their apparent functional discrepancies primarily stem from divergent evolution of their gene regulatory regions, rather than of their respective biochemical properties. Notably, we definitively establish that neither TF directly specifies specific somatosensory neuron identities. This result, combined with : (i) a new series of birth-dating experiments supporting that most mechano/proprioceptive and thermo/nociceptive subtypes are predominantly born during successive time-windows; and (ii) analyses of complementary transgenic mouse models in which opposite temporal shifts of neurogenesis -either delayed onset or premature arrest-have predictable opposite impacts on their DRG content, highlight that the fate choice of somatosensory precursors toward the mechano/proprioceptive or thermo/nociceptive lineages critically depends on the timing of their birth.

## Introduction

During development, the multiple neuronal types that form the central (CNS) and peripheral (PNS) nervous systems are generated at specific times and locations and in definite numbers from neural stem cells, notably through the coordinated action of several families of transcriptional regulators. Among them, the proneural basic-helix-loop-helix (bHLH) family of transcription factors (TFs) plays a pivotal role by controlling neurogenesis and influencing neuronal fate specification (Bertrand et al., 2002; Huang et al., 2014). In Vertebrates, many bHLH TFs have been identified, but only a subset has a clear proneural activity in vivo. Those notably include Mash1/Ascl1 of the Ascl subfamily, Atoh1 and Atoh7 of the Atoh subfamily or Neurog1 and Neurog2 of the Neurogenin subfamily (Huang et al., 2014; Baker and Brown, 2018). Members of different subfamilies generally control neurogenesis in complementary regions of the CNS and PNS and participate in the specification of distinct neuronal types (Huang et al. 2014). Accordingly, gene swapping strategies in mice have highlighted their functional divergences (Baker and Brown, 2018), as shown for *Atoh1*and *Neurog1* (Jahan et al., 2015), *Ascl1* and *Atoh7* (Hufnagel et al., 2013) or *Ascl1* and *Neurog2* (Fode et al., 2000; Parras et al., 2002; Pattyn et al., 2004, 2006; Kele et al. 2006, McNay et al; 2006). However, such approaches have not been employed to assess to which extent paralogous genes of the same subfamily are functionally interchangeable. Replacement of drosophila *atonal* by its mouse orthologues *Atoh1* or *Atoh7* has nevertheless provided indirect evidence that both paralogous TFs are not fully equivalent (Weinberger et al., 2017). In contrast, this question remains largely open for the Neurogenin subfamily. Neurog1 and Neurog2 share a close, though not identical, bHLH domain, and have largely dissimilar NH2- and COOH-terminal regions (Hand et al., 2005; Figure S2A). With few exceptions (e.g Anderson et al., 2006; Kele et al., 2006), they are often detected in the same regions of the developing nervous system, albeit with distinct spatio-temporal patterns, where they fulfill specific, complementary and/or partially redundant roles (Sommer et al., 1996; Fode et al., 1998, 2000; Ma et al., 1998, 1999; Scardigli et al., 2001; Gowan et al., 2001; Hand et al., 2005; Ma et al., 2008; Shaker et al., 2012; Takano-Maruyama et al., 2012; Dixit et al., 2014; Han et al., 2018; Venteo et al., 2019). The developing dorsal root ganglia (DRG) provide a good model illustrating their dynamic expression and complementary requirements. DRG contain a wide variety of specialized neurons involved in the detection of mechanical, proprioceptive, thermal or nociceptive environmental stimuli. Based on anatomical, functional and developmental properties, somatosensory neurons are commonly classified into the mechano/proprioceptive and thermo/nociceptive classes. Both classes ultimately comprise multiple functional subtypes, and it is well-documented that this intra-lineage diversity is progressively established until postnatal stages through binary fate decisions governed by the interplay of environmental cues and intrinsic transcriptional modules (Levanon et al., 2002; Kramer et al., 2006; Chen et al., 2006; Abdo et al., 2011; Scott et al., 2011; Lallemend and Ernfors, 2012; Usoskin et al., 2015; Faure et al., 2020; Sharma et al., 2020; Cranfill & Luo, 2021; Meltzer et al., 2021; de Nooj, 2022). In contrast, the timing and mechanisms by which the mechano/proprioceptive and thermo/nociceptive lineages initially segregate remain elusive and debated (e.g. Sharma et al., 2020; Faure et al, 2020). All somatosensory neurons are produced within a fixed embryonic period, between E9.5 and E13.5, from heterogeneous neural crest (NC)-derived progenitors that emigrate from the dorsal neural tube as “early” or “late” populations, with limited or extensive proliferative capacities, respectively (Serbedjiza et al., 1990; Frank and Sanes, 1991; Maro et al., 2004; Marmigere and Ernfors, 2007; George et al., 2010; Gresset et al., 2015; Ohayon et al., 2015; Vermeiren et al., 2020). These progenitor pools differentiate during two main successive waves of neurogenesis classically assumed to contribute to specific neuronal amounts and identities: the first produces 20% of the DRG neurons between E9.5 and E11.5, mainly mechano/proprioceptors; the second generates 80% of the DRG neurons between E11.5 and E14, mostly thermo/nociceptors (Figure S1A; Lawson and Biscoe, 1979; Kitao et al., 1996; Ma et al., 1999; Lallemend and Ernfors, 2012; Ohayon et al., 2015; Vermeiren et al., 2020). However, recent studies have partly challenged this classical view by suggesting that both functional classes of neurons exhibit more extensive overlapping birth periods and segregate at later stages than previously thought (Sharma et al., 2020; Landy et al., 2021; Meltzer et al., 2021). In this system, *Neurog1* and *Neurog2* exhibit dynamic and complementary expression profiles which roughly correlate with the two neurogenic waves. *Neurog2* is the first to be induced from E9, in all migrating NC-derived somatosensory progenitors at some point, and is subsequently maintained in the coalescing DRG until E11.5, specifically during the first neurogenic period (Figure S1A). *Neurog1* is induced slightly later, at E9.5, under the transient control of *Neurog2*, in post-migratory progenitors settling in the DRG anlagen, and is maintained until E14, encompassing the second neurogenic period (Figure S1A; Ma et al., 1999; Venteo et al., 2019; Soldatov et al., 2019; Faure et al., 2020; Sharma et al., 2020). Accordingly, in *Neurog1^-/-^*knock-outs (KO), the second wave of neurogenesis is specifically abolished and most thermo/nociceptors are missing (Figure S1B; Ma et al., 1999; Bachy et al., 2011; Ohayon et al., 2015). In addition, the complete agenesis of all neuronal classes in *Neurog1^-/-^;Neurog2^-/-^* double-knockouts (dKO) indicates that *Neurog2* control the first neurogenic wave and the formation of mechano/proprioceptors (Figure S1C; Ma et al., 1999). Nevertheless, the individual requirement of *Neurog2* is more complex (Figure S1D). In *Neurog2^-/-^* single-KO, *Neurog1* expression and onset of neurogenesis are delayed by 1.5 days, and during this transient “dKO-like” state, pools of progenitors either die or switch fate to become melanoblasts. After E10.5, however, *Neurog1* is eventually induced in these KO and neurogenesis is in turn initiated, albeit at a lower rate due to progenitor depletion, resulting in a global neuronal deficit of about 30% affecting all functional classes (Figure S1D; Ma et al., 1999; Venteo et al., 2019). Therefore, during DRG development *Neurog1* and *Neurog2* play unique, complementary and only partially redundant roles in maintaining the integrity of somatosensory progenitors and ensuring the generation of accurate neuronal numbers and identities. Notably, Neurog2 seems the sole factor responsible for the repression of the melanocyte fate, while Neurog1 appears the only factor involved in the specification of the thermo/nociceptive identity. However, it remains unclear whether these apparent functional discrepancies reflect distinct biochemical properties between the two TFs and/or their unique spatio-temporal expression patterns. Here, we have assessed the functional equivalence of *Neurog1* and *Neurog2* in the developing DRG using a classical in vivo gene replacement approach. Our findings definitively establish that, despite subtle divergences masked by gene-dosage compensation, *Neurog1* and *Neurog2* are essentially functionally interchangeable in all aspects of DRG development, including fate specification, indicating that their functional specificities primarily reflect their complementary expression patterns. This result, along with new birth dating studies and functional analyses on complementary transgenic mice models in which defined alterations of the timing and pace of neurogenesis have predictable consequences on their DRG content, underscores that the identity of somatosensory precursors as either mechano/proprioceptor or thermo/nociceptor, is early determined but critically relies on the timing of their differentiation.

## Materials and Methods

### Animal models

*Neurog1^2neo^* (*1^2neo^*) and *Neurog2^1neo^*(*2^1neo^*) knock-in (KI) lines were first produced via a classical homologous recombination-based strategy in embryonic stem (ES) cells, using specific targeting constructs (Figures S2C,F). To generate the *1^2neo^*targeting construct, the so-called PBS-X7.5 and PBS-E8.0 *Neurog1* genomic clones covering the entire region of the *Neurog1* locus were used (gift of Dr. Q. Ma). The 5’-arm of homology consisted in a 3.56kb fragment ranging from an upstream XhoI site to the start codon of the endogenous *Neurog1* coding sequence (CDS). The region of the start codon was slightly modified to insert a NsiI restriction site. The CDS of *Neurog2* was then amplified by PCR and cloned into the NsiI site. An “*ires-gfp*” reporter cassette and a “*floxed-pgk-neo*” positive selection cassette (respective gifts of Drs J. Livet and J-F. Brunet) were then introduced. The 3’-arm of homology consisted in a 3.15kb fragment ranging from the endogenous stop codon of *Neurog1* to a downstream XhoI site. Finally, a “*pgk-DTA*” negative selection cassette was introduced (Figure S2C). To generate the *2^1neo^* targeting construct, the so-called PBS-Λ7A genomic clone covering the *Neurog2* locus was used (gift of Dr Q. Ma). The 5’-arm of homology consisted in a 3.15kb fragment ranging from an upstream HindIII site to the start codon of the endogenous *Neurog2* CDS. The region of the start codon was again modified to insert a NsiI restriction site. The CDS of *Neurog1* was then amplified by PCR and cloned into the NsiI site. The “*<u>ires-gfp</u>*” and the “*floxed-pgk-neo*” cassettes were also introduced. The 3’- arm of homology consisted in a 4.5kb fragment ranging from a XbaI site encompassing the stop codon of the endogenous *Neurog2* CDS to a downstream NsiI site. Finally, the “*pgk- DTA*” negative selection cassette was also inserted (Figure S2F). Both constructs were sequence-verified (Cogenics). ES cells were then electroporated, grown and selected using Geneticin (G418; Sigma), and resistant clones were tested for accurate homologous recombination events by Southern blot analyses. For the *1^2neo^* allele, genomic DNA was digested by XbaI, migrated in a 0.8% agarose gel, blotted onto a Nylon membrane (Hybond N+, Amersham) and hybridized with a P^32^-labeled DNA probe corresponding to a XbaI/XhoI fragment located just upstream of the 5’-arm of homology (Figure S2C). Membranes were then analyzed using a phosphoimager. The size of the WT allele was of 11kb while the size of the KI allele was around 6kb (Figure S2D). For the *2^1neo^* allele, genomic DNA was digested by NsiI and a DNA probe corresponding to a NsiI/HindIII fragment upstream of the 5’-arm of homology was used (Figure S2F). The size of the WT allele was of 5.65kb, and that of the KI allele was of 3.95kb (Figure S2G). Recombinant ES clones were then used to generate chimeric males (S.E.A.T facility, Villejuif) that were subsequently bred with WT C57BL/6J females to stably establish the *1^2neo^* and *2^1neo^* transgenic mouse strains which both still contained the “*floxed-pgk-neo*” cassette. *1^2neo^* and *2^1neo^* animals were genotyped by PCR. For *1^2neo^* mice, the following pairs of oligonucleotides were used: i) Neurog1KI2F (5’- acggacagtaagtgcgct-3’) and Neurog2KI1R (5’-ttggctttgacgcgacgact-3’) to assess the *Neurog1^WT^* allele; ii) Neurog1KI2F and Neurog1KI2R (5’-tcgtgagcttggcatcct-3’) for the *Neurog1^2neo^* allele; and iii) NeoF (5’-ctattcggctatgactgggc-3’) and NeoR (5’- aatatcacgggtagccaacg-3’) for the *neomycin* gene. Expected band sizes were respectively of 368bp, 526bp and 630bp (Figure S2E). For *2^1neo^* animals: i) VD187 (5’-ggacattcccggacacacac- 3’) and Neurog1KI2R oligonucleotides were used to assess the *Neurog2^WT^*allele; ii) Neurog2KIF (5’-agcgacactttaccctgtcc-3’) and Neurog2KI1R for the *Neurog2^1neo^* allele; and iii) NeoF and NeoR for the *neomycin* gene. Expected band sizes were respectively of 779bp, 361bp and 630bp (Figure S2H). To excise the “*floxed-pgk-neo*” cassette, the *CMV-Cre* ubiquitous deleter mice (Schwenk et al., 1995) was used, thus allowing the production of the *Neurog1^2^* and *Neurog2^1^* mice that represent “clean” KI (Figures S2C,F). These animals were PCR-tested for the deletion of the *neomycin* gene and were subsequently genotyped using the same pairs of oligonucleotides as described above (Figures S2E,H). Note that both KI strains are viable and fertile at the homozygous state.

*Neurog1* (Ma et al., 1999) and *Neurog2* (Andersson et al., 2006) knockouts (KO) lines were also used. They were maintained and genotyped as previously described (Venteo et al., 2019). *1^2neo/2neo^*;*Neurog2^-/-^*, and *2^1neo/1neo^*;*Neurog1^-/-^*compound transgenic embryos as well as *Neurog1^2/2^*;*Neurog2^-/-^*, and *Neurog2^1neo/1neo^*;*Neurog1^-/-^* KI;KO models were obtained by successive breeding of appropriate KI and KO animals. Histological analyses were performed on embryos issued from single-KI, single-KO or compound KI;KO lines harvested between embryonic day (E) 9.5 and E18.5. The morning of the vaginal plug was considered as embryonic stage E0.5. Parents were provided ad libitum with standard mouse pellet food and water and housed at room temperature with a 12h light/dark cycle. All procedures were conducted according to the French Ministry of Agriculture and the European Community Council Directive no. 86/609/EEC, OJL 358. Protocols were validated by the Direction Départementale des Services Vétérinaires de l’Hérault (Certificate of Animal Experimentation no. 34-376).

### B16F10 cell cultures and transfection of expression vectors

B16F10 cells (ATCC) were cultured in DMEM/F12 medium (Lonza) supplemented with 10% heat-inactivated fetal bovine serum (FBS, Gibco) and Penicillin/Streptomycin (Gibco). Cell line were maintained at 37°C in a humified incubator with 5% CO_2_. B16F10 cells were transfected at 80% confluency in 6-well plates using lipofectamine 2000 (Thermo Fisher Scientific), Opti-MEM Reduced Serum Medium (Gibco) and 1µg of total plasmid DNA. Cells were fixed with 4% paraformaldehyde for 15 min at room temperature 24 hours post-transfection. GFP, Neurog1, Neurog2, Neurog1-bhlh2 and Neurog2-bhlh1 were expressed from the CMV promoter of the pCS2+ vector. Myc-mNeurog1-PCS2 was kindly provided by Eric Bellefroid (ULB, Belgium). HA-mNeurog2-PCS2 was cloned using the In-Fusion cloning strategy (Takara) using PCR product generated with Phusion DNA polymerase (NEB). The primer sequences used were: Fwd:5’- CTTTTTGCAGGATCCCATCGATGCCACCATGTATCCGTATGATGTTCCGGATTATG CATTCGTCAAATCTGAGACTCTGGAGTTG-3’ and Reverse: 5’- CTATAGTTCTAGAGGCTCGAGCTAGATACAGTCCCTGGCGAGG-3’. Chimeras Neurog1-bHLH2 and Neurog2-bHLH1 were synthetized using the Genewiz Gene synthesis service. All constructs were confirmed by Sanger sequencing.

### Quantitative RT-PCR

RNA extractions were performed using RNeasy Mini kit (Qiagen) according to the manufacturer’s instructions. A total of 1µg of RNA was used to retrotranscribe in cDNA with the Superscript II First-Strand Synthesis System (Thermo Fhisher Scientific). Real-time PCR was carried out using SYBR Green I dye detection on the Light Cycler system (Roche Molecular Biochemicals). The results are expressed as relative mRNA levels of gene expression and obtained using the ΔΔCT-method. The normalization was performed with GAPDH as a housekeeping gene and compared to control condition. The primer sequences used were: *GAPDH* Fwd: 5’-GTGAAGGTCGGTGTGAACG-3’; *GAPDH* Rev: 5’-AGGGGTCGTTGATGGCAACA-3’; *MitF* Fwd: 5’- ACCTCTGAAGAGCAGCAGTT-3’; *Mitf* Rev: 5’- AGTTGCTGGCGTAGCAAGAT-3’

### Immunofluorescence staining

Primary antibodies used in this study were as follows: Chicken anti-GFP (1/1000; Abcam), Goat anti-TrkB (1/2000; R&D Systems), Goat anti-TrkC (1/1000; R&D Systems), Mouse anti-MitF (1/100; C5/D5 Sigma), Rabbit anti-activated Caspase3 (1/2000; Cell Signaling), Rabbit anti-Parvalbumin (PV; 1/10000; SWANT) and Rabbit anti-TrkA (1/500; Millipore). Secondary antibodies (Molecular Probes) used were as follows; Donkey anti-rabbit IgG Alexa Fluor 594-conjugated (1/2000); Donkey anti-goat IgG Alexa Fluor 488-conjugated (1/1000); Goat anti-chicken IgG Alexa Fluor 488-conjugated (1/1000). For staining on cryosections, slides were dried 20 min at RT, rinsed with PBS, blocked for 30 min at RT in 4% Serum-PBS-0.1% Triton X100 and incubated o.n. at 4°C with the appropriate primary antibody(ies) diluted in the same buffer. Slides were then washes 3 times 10 min in PBS-0.1% Triton X100 and incubated 1h at RT with secondary antibody(ies) diluted in the same buffer. They were then washed 3 times 10 min at RT, mounted under coverslips in Mowiol and rapidly imaged. Note that pictures showing ISH or immunofluorescence staining are representative of at least three independent animals.

For staining on cell cultures, cells were permeabilized in PBTT (PBS 0.1% Tween-20; 0.5% triton X-100) and then blocked in 3% BSA-PBS for 45 min before being exposed to primary antibodies for 1h at room temperature. After three PBS washes, cells were incubated with secondary antibodies for 1h at room temperature. Cells were stained for 2 min with Hoechst diluted to 10 µg/mL in PBS.

### In situ hybridization (ISH) on cryosections

Antisense RNA probes for *Cgrp, gfp, MafA*, *Mitf*, *MrgprD*, *NeuroD*, *Neurog1, Neurog2, Ret, SCG10*, *Sox10, TrkA, TrkB, TrkC, TrpM8* and *TrpV1* were labeled using the Dig RNA labeling kit (Roche) and purified on G50 columns (GE Healthcare). For staining on cryosections, staged embryos were dissected out in cold Phosphate Buffer Saline (PBS), fixed overnight (o.n.) at 4°C in 4% Paraformaldehyde(PFA)- PBS, cryoprotected in 20% Sucrose-PBS o.n. at 4°C, embedded in OCT-compound (Tissue- Tek) and stored at -80°C until use. 14µm thoracic transverse sections were cut using a cryostat (Microm), placed on glass slides (Superfrost Plus, Menzel-Gläser) and stored at - 20°C until use. For the staining procedure, slides were dried at room temperature (RT) for 20 minutes (min) and incubated o.n. at 70°C with the appropriate probes diluted 1/100-1/200 in the hybridization buffer (50% Formamide-10% Dextran Sulfate-1X Salt Solution-1mg/ml yeast tRNA-1X Denhardt’s). They were washed twice 1h at 70°C in 50% Formamide-1X Sodium/Sodium Citrate buffer-0.1% Tween 20. They were then washed 3 times 10 min at RT in MABT (1X Maleic acid buffer-0.1% Tween20), blocked in 20% Sheep Serum-2% Blocking Reagent (Roche)-MABT and incubated o.n. at 4°C with the anti-DIG antibody (Roche) diluted 1/2000 in the same buffer. Slides were washed 3 times for 10 min at RT in MABT, incubated twice for 15 min in buffer B3 (100mM Tris pH 9.5, 100mM NaCl, 50mM MgCl2, 0.1% Tween 20) and incubated with the NBT and BCIP substrates (Roche) diluted in B3. Slides were washed several times in water for 2h, dried at RT and mounted in Mowiol under coverslips.

### Birth dating analyses

Birth dating experiments were performed using intraperitoneal injections of BrdU into pregnant mice (2mg/mouse in PBS) at either E9.5, E10.5, E11.5, E12.5 or E13.5. Embryos were then collected at E18.5 and a column containing their spinal cord and DRG was dissected out, fixed in 4% Paraformaldehyde(PFA)-PBS, cryoprotected in 20% Sucrose-PBS and embedded in OCT-compound. 14µm thoracic transverse sections were then processed for double-staining, either by combined ISH and BrdU immunofluorescence (*TrkB*/BrdU, *TrkC*/BrdU, *MafA*/BrdU, *MrgprD*/BrdU, *Cgrp*/BrdU, *TrpM8*/BrdU and *TrpV1*/BrdU) or by double-immunofluorescence (PV/BrdU) (Figure S5A). For ISH and BrdU double-staining, ISH was first performed until appropriate coloration of the cytoplasm. Slides were then washed in PBS and incubated in 10 mM sodium citrate buffer (pH 6.0) at 95°C for 15 min, then at room temperature (RT) for further 15 min, and washed several times in PBS at RT. They were then subjected to immunofluorescence staining to detect BrdU in the nucleus. For double immunofluorescence, slides were dried at RT, washed once in PBS and incubated in 10 mM sodium citrate buffer (pH 6.0) at 95°C for 15 min, then at room temperature (RT) for further 15 min, and washed several times in PBS at RT. They were then subjected to double-immunostaining staining as described above.

### Quantification and statistical analyses

Quantitative analyses on B16F10 cells were performed on 3 independent transfection experiments. Quantitative analyses on mice were carried out on at least 3 independent animals for each genotype or conditions. Thoracic DRG sections processed either for ISH or immunofluorescence were imaged using a Zeiss microscope. Cell counts were performed on 6-8 sections (12-16 hemisections) for each animal, depending on the stage, using the Image J software. Only cells with a clearly identifiable nucleus were counted. For birth dating experiments, only cells with a clear BrdU- positive nucleus was taken into consideration. Nuclei containing only few fluorescent dots were not considered to avoid potential overestimation.

For RT-qPCR experiments, statistical analyses were performed using one-way analysis of variance (ANOVA), followed by a post-hoc Newman-Keuls test. For cell countings in vitro and on sections of mouse embryos, statistical analyses were performed using Unpaired Student *t* tests. Results are expressed as mean ± standard error to the mean (s.e.m). p <0.05 (*), p < 0.01 (**) and p < 0.001 (***) were considered as statistically significant. Note that no statistical methods were used to predetermine sample sizes, but numbers of animals and sections analyzed for each genotype and for each stage are consistent with those typically used in the field.

## Results

### Generation and Validation of *Neurog1* and *Neurog2* Knock-in Mouse Models

To assess functional similarities and divergences of *Neurog1* and *Neurog2* in the developing DRG, we generated knock-in (KI) mouse lines where their coding sequences (CDS) were mutually swapped, while preserving the overall genes structure. This was facilitated by the relatively simple organization of their respective loci in which the CDS is encoded by a single exon (Figures S2C,F). Using conventional gene targeting in mouse embryonic stem (ES) cells, the endogenous *Neurog1-CDS* and *Neurog2-CDS* were precisely replaced by a “*Neurog2-CDS/ires-gfp/Floxed-PGK-neo*” construct (Figure S2C) and a “*Neurog1-CDS/ires- gfp/Floxed-PGK-Neo*” construct (Figure S2F), respectively. After ES cells electroporation, selection and genotyping, we identified several recombinant clones (Figures S2D,G) that were used to initially produce the so-called *Neurog2^KINeurog1neo^* (hereafter referred to as *2^1neo^*transgenic line; Figure S2C) and *Neurog1^KINeurog2neo^* (hereafter referred to as *1^2neo^* transgenic line; Figure S2F) lines, which both contain the *Floxed-PGK-neo* cassette. Having in mind that the *PGK* promoter can perturb the activity of the locus in which it is inserted (e.g Scacheri et al, 2001; see below), both lines were subsequently crossed with a *CMV-Cre* deleter line to establish the *Neurog2^KINeurog1^*(hereafter referred to as *Neurog2^1^* or *2^1^* line) and *Neurog1^KINeurog2^* (hereafter referred to as *Neurog1^2^* or *1^2^* line) lines which constitute “clean” KI models (Figures S2C,E,F,H; see below). Each model was then validated by analyzing the expression of the different transgenes, focusing on the developing DRG. For the *Neurog2^1/1^* KI animals, considering the transient expression of *Neurog2* in this structure between E9 and E11.5 and its extensive overlap with *Neurog1* at these stages (Ma et al., 1999; Figures S1A and 1M), we initially took advantage of the *gfp* reporter gene. Thereby, we determined that the spatio-temporal profile and number of *gfp*-positive (+) cells were similar to that of *Neurog2*+ cells in WT, starting at E9 and ending at E11.5 (Figures 1A-C). Consistent with this, at E9.5, the number of *Neurog1*+ cells in *Neurog2^1/1^* embryos was increased compared to WT, and was similar to *Neurog2*+ cells in WT (Left graph in Figure 1C). From E10.5, when endogenous *Neurog1* expression normally overrides that of *Neurog2* in WT, no difference in the number of *Neurog*1+ cells was observed between *Neurog2^1/1^* and WT embryos (Middle and right graphs in Figure 1C). Importantly, the larger amount of *Neurog1*+ cells at E10.5 in *Neurog2^1/1^* compared to *Neurog2*+ cells in WT (Middle graph in Figure 1C), indicates that exogenous Neurog1 is able to activate the endogenous *Neurog1* locus, as Neurog2 normally does (Summarized in Figures 1M,N; Ma et al., 1999; Venteo et al., 2019). In parallel, analyses on *Neurog1^2/2^* embryos between E10.5 and E13.5, revealed that the spatio-temporal profile and number of *Neurog2*+ cells were fully reminiscent of that of *Neurog1*+ cells in WT (Figures 1D-F). Notably, from E11.5 when the *Neurog2* locus is normally turned off in WT (Figure 1A), expression of exogenous *Neurog2* was still expectedly detected in the KI animals, as *Neurog1* in WT (Figures 1D,E; Middle and right graphs in Figure 1F; summarized in Figure 1O). These results thus show that in *Neurog2^1/1^*and *Neurog1^2/2^* KI models, exogenous *Neurog1* and *Neurog2* are properly expressed (Summarized in Figures 1M-O).

**Figure 1.**
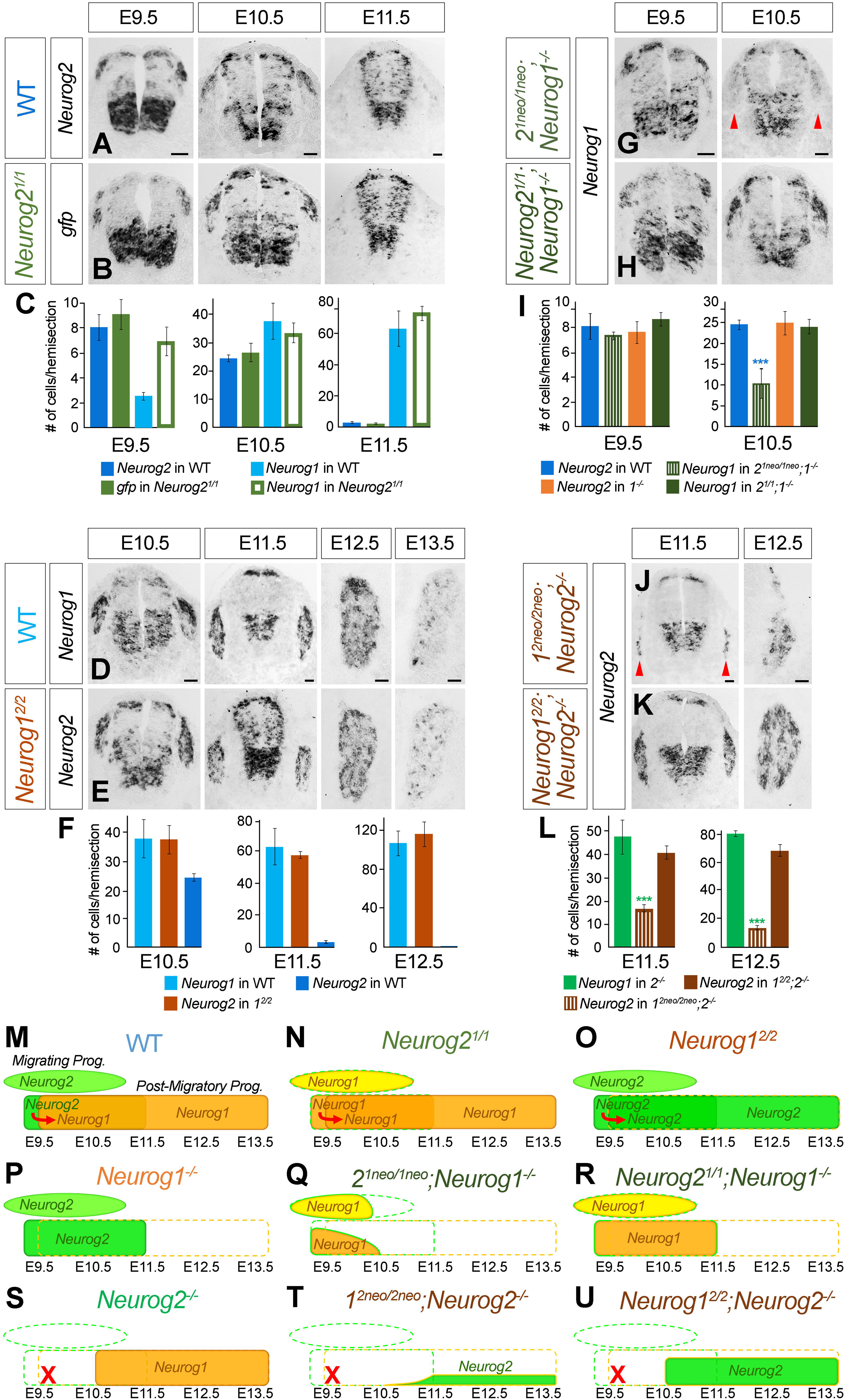
Characterization and validation of Knock-In alleles of *Neurog1* and *Neurog2*. **A,B.** Representative images of *in situ* hybridization for *Neurog2* in WT (A) and for *gfp* in *Neurog2^1/1^* KI (B) embryos on transverse thoracic sections of the spinal cord at E9.5, E10.5 and E11.5, as indicated. Scale bars, 50µm. **C.** Quantitative analysis of the numbers of *Neurog2*+ cells in WT (dark blue columns), of *gfp*+ cells in *Neurog2^1/1^* KI (green columns) and of *Neurog1*+ cells in WT (light blue columns) and *Neurog2^1/1^*(green-framed white columns) KI at E9.5, E10.5 and E11.5, as indicated. Data are represented as means ± SEM. **D,E.** Representative images of *in situ* hybridization for *Neurog1* in WT (D) and for *Neurog2* in *Neurog1^2/2^* (E) on transverse thoracic spinal cord or DRG sections at E10.5, E11.5, E12.5 and E13.5, as indicated. Scale bars, 50µm. **F.** Quantitative analysis of the numbers of *Neurog1*+ cells in WT DRG (light blue columns), and of *Neurog2*+ cells in *Neurog1^2/2^* (light brown columns) and WT DRG (dark blue columns) at E10.5, E11.5 and E12.5 as indicated. Data are represented as means ± SEM. **G,H.** Representative images of *in situ* hybridization for *Neurog1* in *2^1neo/1neo^;Neurog1^-/-^* (G) and *Neurog2^1/1^;Neurog1^-/-^* (H) embryos on transverse thoracic spinal cord sections at E9.5 and E10.5, as indicated. Red arrowheads in G point to the coalescing DRG that abnormally express low levels of *Neurog1* in E10.5 *2^1neo/1neo^;Neurog1^-/-^* embryos. Scale bars, 50µm. **I.** Quantitative analysis of the numbers of *Neurog2*+ cells in the DRG of WT (dark blue columns) and *Neurog1^-/-^* KO (orange columns), and of *Neurog1*+ cells in the DRG of *2^1neo/1neo^;Neurog1^-/-^*(striped dark green columns) and *Neurog2^1/1^;Neurog1^-/-^*(dark green columns) embryos at E9.5 and E10.5, as indicated. Data are represented as means ± SEM. ***p < 0.001. Blue asterisks indicate significant differences with WT. **J,K.** Representative images of *in situ* hybridization for *Neurog2* in *1^2neo/2neo^;Neurog2^-/-^* (J) and *Neurog1^2/2^;Neurog2^-/-^* (K) embryos on transverse thoracic spinal cord or DRG sections at E11.5 and E12.5, as indicated. Scale bars, 50µm. **L.** Quantitative analysis of the numbers of *Neurog1*+ cells in the DRG of *Neurog2^-/-^* KO (green columns), and of *Neurog2*+ cells in the DRG of *1^2neo/2neo^;Neurog2^-/-^* DRG (striped-dark brown columns) and *Neurog1^2/2^;Neurog2^-/-^*(dark brown columns) embryos at E11.5 and E12.5, as indicated. Data are represented as means ± SEM. ***p < 0.001. Green asterisks indicate significant differences with *Neurog2^-/-^* KO. **M-U.** Schematic summary of the dynamic expression profiles of *Neurog2* (green colors) and *Neurog1* (orange colors) in migrating (oval frames) and/or post-migratory (rectangle frames) somatosensory progenitors in WT (M), *Neurog2^1/1^* (N), *Neurog1^2/2^* (O), *Neurog1^-/-^*(P), *2^1neo/1neo^;Neurog1^-/-^* (Q), *Neurog2^1/1^;Neurog1^-/-^* (R), *Neurog2^-/-^*(S), *1^2neo/2neo^;Neurog2^-/-^* (T) and *Neurog1^2/2^;Neurog2^-/-^* (U) embryos between E9.5 and E13.5. Red arrows in M,N,O illustrate the ability of earlier-expressed gene from the *Neurog2* locus in regulating the onset of the later-expressed gene from the *Neurog1* locus. Red crosses illustrate that when the endogenous *Neurog2* locus is deleted, expression onset from the endogenous *Neurog1* locus is impaired. Note that the PGK-neo cassette in the *2^1neo/1neo^;Neurog1^-/-^* (Q) and the *1^2neo/2neo^;Neurog2^-/-^* (T) models affects the activities of the *Neurog2* and *Neurog1* loci, respectively.

During the course of this study, we noticed that in the *2^1neo^*and *1^2neo^* transgenic lines, the *PGK-neo* cassette perturbs the expression of the knocked-in genes. This effect was particularly evident in *2^1neo/1neo^* embryos, where the *gfp* reporter gene was induced at E9.5 but prematurely down-regulated by E10.5, one day earlier than *Neurog2* in WT (Figures S3A,B). Surprisingly though, the number of *Neurog1*+ cells was essentially normal in these animals over time, including at E10.5 (Figures S3C,D). This likely reflects that transient expression of exogenous *Neurog1* is sufficient to activate the endogenous *Neurog1* locus which, in turn, maintains its own expression (Summarized in Figure S3E). Supporting this, analyses on *2^1neo/1neo^*;*Neurog1^-/-^*compound transgenic embryos in which the endogenous *Neurog1* locus was additionally deleted, revealed that *Neurog1* solely expressed from the *Neurog2* locus was induced at E9.5, but became barely detectable by E10.5 (Figures 1G,I; summarized in Figure 1Q). Importantly, this premature down-regulation of the *Neurog2* locus was not observed in *Neurog1^-/-^* single-KO or in *Neurog2^1/1^*;*Neurog1^-/-^* KI/KO animals (Figures 1H,I; summarized in Figures 1P,R). In the same line, analyses of *1^2neo/2neo^;Neurog2^-/-^* compound transgenic embryos revealed that induction of exogenous *Neurog2* from the *Neurog1* locus was further delayed, and eventually occurred in fewer cells, compared to the endogenous *Neurog1* in *Neurog2^-/-^* single-KO (Figures 1J,L; summarized in Figures 1S,T; Ma et al., 1999; Venteo et al., 2019). Importantly, these defects were not observed in *Neurog1^2/2^*;*Neurog2^-/-^*KI/KO embryos, although we noted a slight tendency towards a reduction in the number of *Neurog2*+ cells compared to that of *Neurog1*+ cells in *Neurog2^-/-^* single-KO (Figure 1K,L; summarized in Figures 1S,U; see below). These data thus establish that presence of the “*PGK-neo* cassette” shortens the periods of activity of the *Neurog2 and Neurog1* loci, albeit in strikingly opposite ways: in *2^1neo/1neo^*;*Neurog1^-/-^* embryos, it causes a premature down-regulation of the *Neurog2* locus compared to WT or *Neurog1^-/-^* single-KO (Figures 1P,Q), while in *1^2neo/2neo^;Neurog2^-/-^* animal, it triggers a delayed activation of the *Neurog1* locus compared to *Neurog2^-/-^* single-KO (Figures 1S,T).

Altogether, these data validate the *Neurog2^1/1^* and *Neurog1^2/2^* single-KI lines, as well as the *Neurog2^1/1^;Neurog1^-/-^* and *Neurog1^2/2^;Neurog2^-/-^* KI/KO lines, as reliable and complementary models for evaluating the functional equivalence of *Neurog1* and *Neurog2* in the developing DRG (Figures 1M,N,O,R,U). They also fortuitously establish the *2^1neo/1neo^*;*Neurog1^-/-^* and the *1^2neo/2neo^;Neurog2^-/-^*compound transgenic lines as interesting tools for studying the consequences of temporal alterations of proneural genes expression in this structure, complementary to the *Neurog1^-/-^* and *Neurog2^-/-^* single-KO models (Figures 1P,Q,S,T).

### Proper *Neurog2* locus activity, regardless of whether it encodes Neurog2 or Neurog1, is essential to repress the melanocyte fate in migrating somatosensory progenitors

We next assessed the extent to which the defects observed in the developing DRG of *Neurog2^-/-^* and *Neurog1^-/-^* KO models are rescued in *Neurog2^1/1^* and *Neurog1^2/2^* KI models, respectively. We first focused on the specific requirement of *Neurog2* in controlling survival and lineage commitment of NC-derived somatosensory progenitors at early developmental stages (Figure S1D; Ma et al., 1999; Soldatov et al., 2019; Venteo et al., 2019). To evaluate whether *Neurog1* can replace *Neurog2* in these processes, we performed a comparative time- course analysis of the *Sox10*+ somatosensory progenitor population between WT, *Neurog2^-/-^* KO and *Neurog2^1/1^* KI animals, from E10.5 to E12.5. In line with previous findings, at E10.5, *Neurog2^-/-^* KO exhibited an increased number of *Sox10*+ cells compared to WT (Figure 2A), due to delayed neurogenesis (Venteo et al., 2019; see also Figures 3A-C). However, from E11.5, this cell population was significantly depleted in these mutants (Figures 2A,B,B’), reflecting that pools of progenitors either undergo apoptosis (Figures 2C,C’) or switch fate toward the melanocyte lineage (Figures 2D,E; quantified in Figure 2N; Venteo et al., 2019). Analysis of *Neurog2^1/1^*KI embryos showed that the number of *Sox10*+ progenitors in their DRG anlagen was strikingly similar to that of WT at all stages analyzed (Figures 2A,B,B’’). Consistent with this, no apoptotic figures at E10.75 (Figure 2C’’) and no supernumerary *Mitf*+ melanoblasts at E11.5 (Figures 2F,N) were detected in these KI animals. These results thus show that *Neurog1* can substitute for *Neurog2* to preserve the integrity of somatosensory progenitors at early stages and, notably, that *Neurog1* has the potential to inhibit the melanocyte fate, even though it is normally not necessary during this process (Venteo et al., 2019). In addition, analysis of *Neurog2^1/1^*;*Neurog1^-/-^*KI;KO embryos established that only two copies of the *Neurog1-CDS* exclusively expressed from the *Neurog2* locus are sufficient to prevent the formation of supernumerary melanoblasts (Figures 2G,N). In contrast, in *Neurog1^2/2^*;*Neurog2^-/-^*KI;KO embryos, re-expressing two copies of the *Neurog2-CDS* from the *Neurog1* locus did not rescue the fate switch phenotype triggered by the loss of *Neurog2* function (Figures 2E,H,N). These data thus show that, regardless of whether Neurog1 or Neurog2 protein is encoded, the specific activity of the *Neurog2* locus is essential for inhibiting the melanocyte fate in a pool of somatosensory progenitors, while that of the *Neurog1* locus is dispensable.

**Figure 2.**
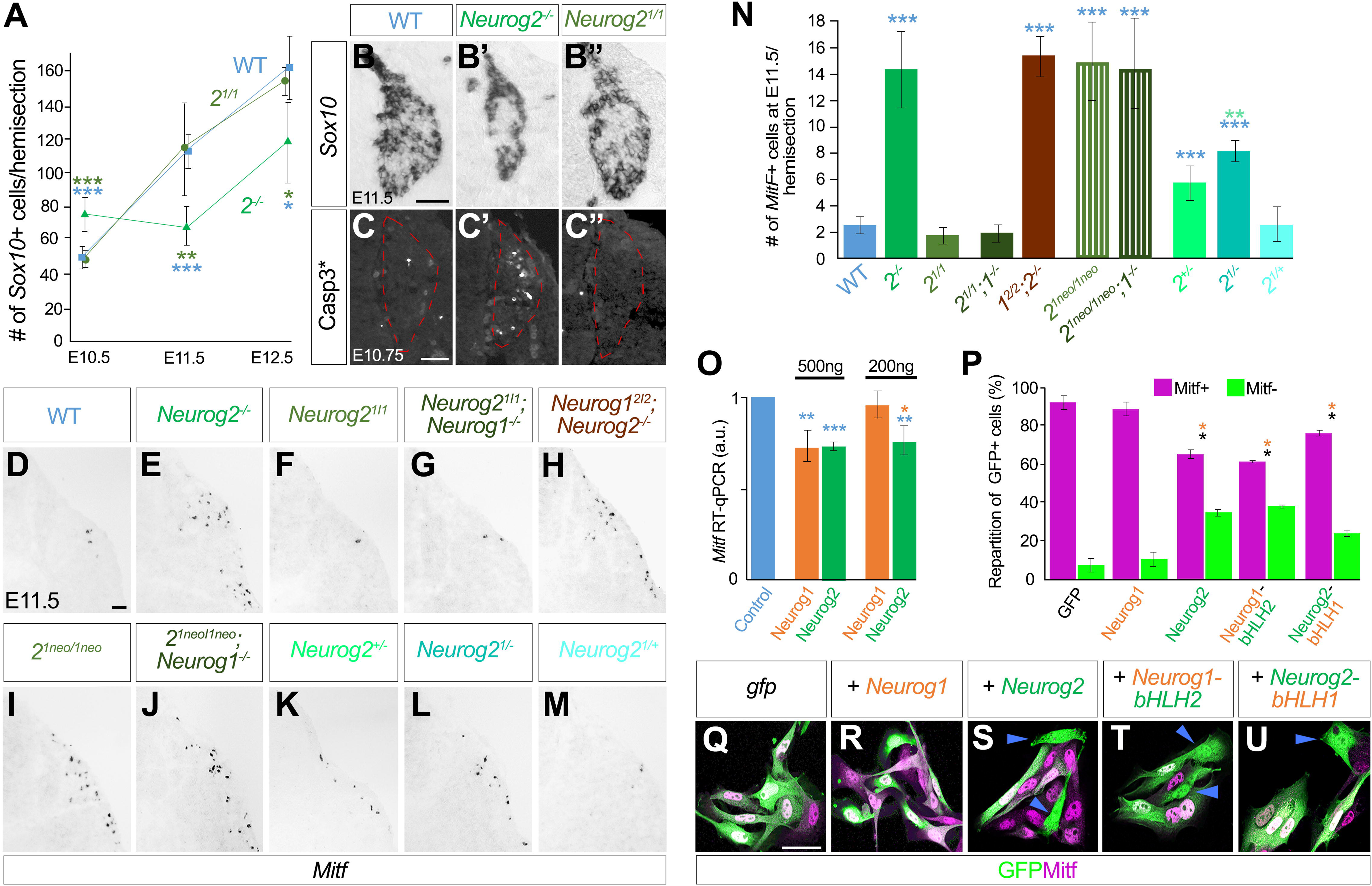
*Neurog1* can replace *Neurog2* function to maintain somatosensory progenitors integrity and repress the melanocyte fate, albeit with slightly reduced efficiency. **A.** Comparative quantitative time-course analysis of the number of *Sox10*+ somatosensory progenitors in WT, *Neurog2^-/-^* (*2^-/-^*) and *Neurog2^1/1^* (*2^1/1^*) embryos between E10.5 and E12.5. Data are represented as means ± SEM. *p < 0.05; **p< 0.01; ***p < 0.001. **B-C’’.** Representative images of *in situ* hybridization for *Sox10* at E11.5 (B-B’’) or of immunofluorescent staining using an activated-Caspase3 (Caps3*) antibody at E10.75 (C-C’’) on transverse thoracic DRG sections of WT (B,C), *Neurog2^-/-^* (B’,C’) and *Neurog2^1/1^*(B’’,C’’) embryos. Red dotted lines in C-C’’ delimit the DRG. Scale bars, 50µm. **D-M.** Representative images of *in situ* hybridization for *Mitf* at E11.5 on transverse thoracic sections of WT (D), *Neurog2^-/-^*(E), *Neurog2^1/1^* (F), *Neurog2^1/1^*;*Neurog1^-/-^*(G), *Neurog1^2/2^*; *Neurog2^-/-^* (H), *2^neo1/neo1^*(I), *2^neo1/neo1^*;*Neurog1^-/-^* (J), *Neurog2^+/-^*(K), *Neurog2^1/-^* (L) and *Neurog2^1/+^* (M) embryos, showing the dorso-lateral migratory path of NCC. Scale bar, 50µm. **N.** Quantification of the number of *Mitf*+ melanoblasts located in the NCC dorso-lateral migratory path of WT, *Neurog2^-/-^* (*2^-/-^*), *Neurog2^1/1^* (*2^1/1^*), *Neurog2^1/1^;Neurog1^-/-^*(*2^1/1^;1^-/-^*), *Neurog1^2/2^*;*Neurog2^-/-^ (1^2/2^*;*2^-/-^*), 2*^1neo/1neo^*, 2*^1neo1/1neo^*;*Neurog1*^-^*^/-^* (2*^1neo1/1neo^*;1*^-/^*^-^), *Neurog2^+/-^* (*2^+/-^*), *Neurog2^1/-^* (*2^1/-^*) and *Neurog2^1/+^*(*2^1/+^*) embryos at E11.5. Data are represented as means ± SEM. *p < 0.05; ***p < 0.001. Blue asterisks indicate significant differences with WT. Light green asterisk indicate significant differences between *Neurog2^+/-^* and *Neurog2^1/-^* embryos. **O.** Quantitative analysis of the level of *Mitf* expression assessed by RT-qPCR on B16F10 cell cultures after transfection of 1µg of a *gfp*-expression vector (blue columns) or after co- transfection of either 500ng or 200ng of a *Neurog1*-expression vector (orange columns) or a *Neurog2*-expression vector (green column) together with either 500ng or 800ng of the *gfp*- expression vector, respectively. Data are represented as means ± SEM. *p < 0.05; **p< 0.01; ***p < 0.001. Blue asterisks indicate significant differences with the control *gfp* condition, while the orange asterisk indicate significant differences between the *Neurog1* and the *Neurog2* conditions at 200ng. **P.** Quantification of the proportion of GFP+Mitf+ (Magenta columns) and GFP+Mitf- (Green columns) B16F10 cells after transfection of 1µg of a control GFP-expression vector, or after co-transfection of 800ng of the gfp-expression vector together with 200ng of expression vectors encoding either native Neurog1, native Neurog2, chimeric Neurog1-bHLH2 or chimeric Neurog2-bHLH1 proteins, as indicated. Data are represented as means ± SEM. *p < 0.05. Black asterisks indicate significant differences with the control GFP condition, while orange asterisks indicate significant differences with the “native Neurog1” condition. **Q-U.** Representative images of co-immunofluorescent staining using GFP (green) and Mitf (magenta) antibodies on B16F10 cells after transfection of control *gfp* vector alone (Q) or co- transfection of GFP and with expression vectors for either native Neurog1 (R), native Neurog2 (S), chimeric Neurog1-bHLHNeurog2 (T) or chimeric Neurog2-bHLHNeurog1 (U) proteins, as indicated on the graph. Blue arrowheads in S, T and U, point to GFP+Mitf- cells. Scale bar, 20µm.

**Figure 3.**
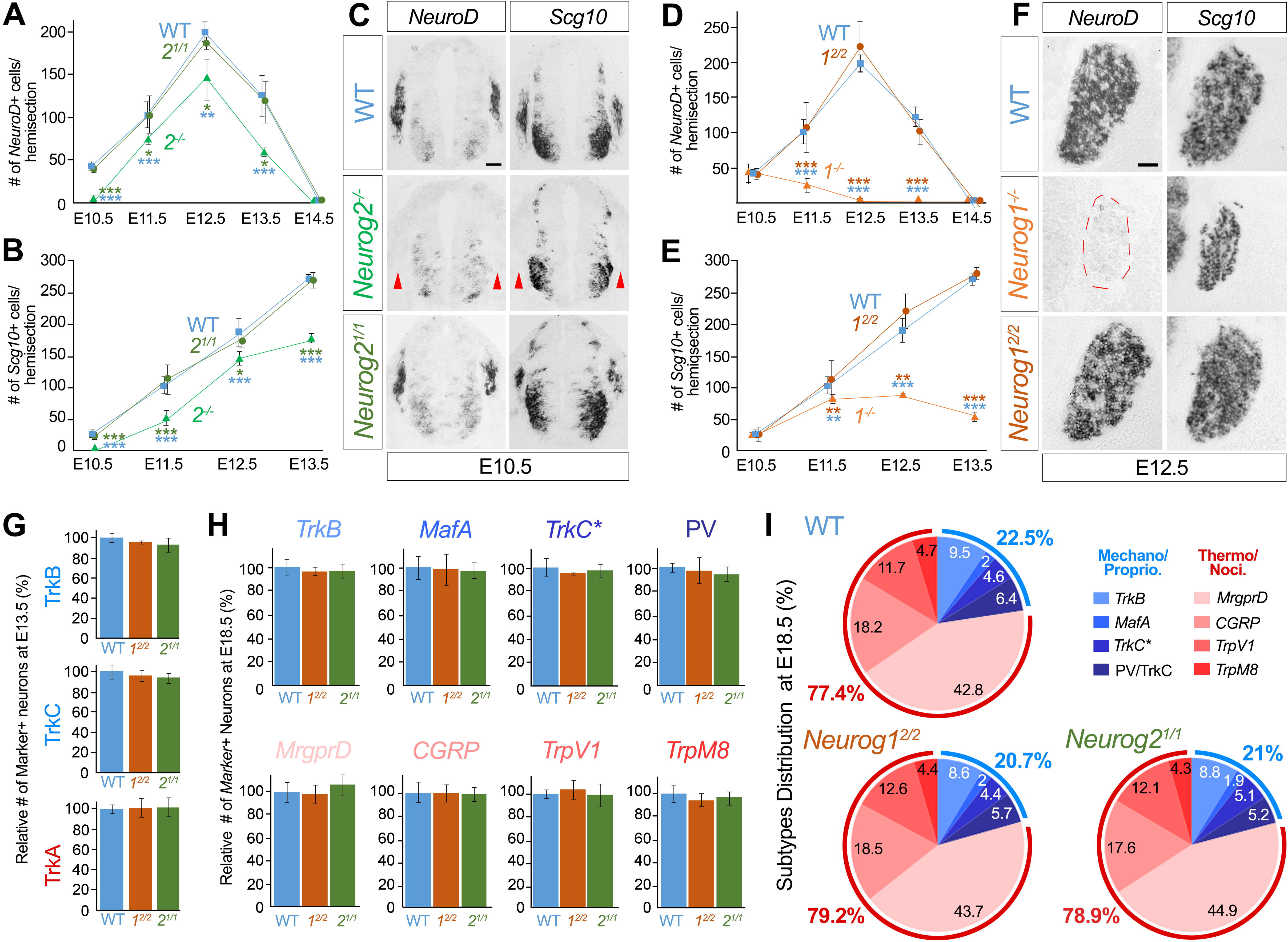
*Neurog1* and *Neurog2* are functionally interchangeable for controlling neurogenesis and fate specification in the developing DRG. **A,B.** Quantitative time-course analysis of *NeuroD*+ differentiating precursors between E10.5 and E14.5 (A) or of *Scg10*+ post-mitotic neurons between E10.5 and 13.5 (B) in WT (blue curves), *Neurog2^-/-^* KO (*2^-/-^*; green curves) and *Neurog2^1/1^* (*2^1/1^*; dark green curves). Data are represented as means ± SEM. *p < 0.05; **p < 0.01; ***p < 0.001. Blue asterisks indicate significant differences with WT. Green asterisks indicate significant differences with *2^1/1^* KI. **C.** Representative images of *in situ* hybridization for *NeuroD* and *SCG10* on transverse spinal cord sections of WT, *Neurog2^-/-^* and *Neurog2^1/1^* embryos at E10.5. Red arrowheads point to the coalescing DRG of *Neurog2^-/-^* KO which are devoid of staining at this stage. Scale bar, 50µm. **D,E.** Quantitative time-course analysis of *NeuroD*+ differentiating precursors between E10.5 and E14.5 (D) and of *Scg10*+ post-mitotic neurons between E10.5 and E13.5 (E) in WT (blue curves), *Neurog1^-/-^* KO (*1^-/-^*; orange curves) and *Neurog1^2/2^* (*1^2/2^*; brown curves). Data are represented as means ± SEM. **p < 0.01; ***p < 0.001. Blue asterisks indicate significant differences with WT. Brown asterisks indicate significant differences with *1^2/2^* KI. **F.** Representative images of *in situ* hybridization for *NeuroD* and *Scg10* on transverse DRG sections of WT, *Neurog1^-/-^* KO and *Neurog1^2/2^* embryos at E12.5. The red dotted line delimits the DRG of *Neurog1^-/-^*KO which are devoid of *NeuroD* staining at this stage. Scale bar, 50µm. **G.** Relative quantification of TrkB+, TrkC+ and TrkA+ neuronal precursors in the DRG of WT (blue columns), *Neurog1^2/2^* KI (*1^2/2^*; brown columns) and *Neurog2^1/1^* (*2^1/1^*, green columns) at E13.5. Data are represented as means ± SEM. **H.** Relative quantification of *TrkB*+, *MafA*+, *TrkC*+/PV- (*TrkC*\*) and PV+ mechano/proprioceptive subtypes, and *MrgprD*+, *Cgrp*+, *TrpV1*+ and *TrpM8*+ thermo/nociceptive subtypes in WT (blue columns), *Neurog1^2/2^* KI (*1^2/2^*; brown columns) and *Neurog2^1/1^* (*2^1/1^*, green columns) at E18.5. Data are represented as means ± SEM. **I.** Graphic summaries showing the distribution of each neuronal subtype analyzed in this study: the *TrkB*+, *MafA*+, *TrkC*+/PV- (*TrkC*\*) and PV+ mechano/proprioceptors (blue colors) and the *MrgprD*+, *CGRP*+, *TrpV1*+ and *TpM8*+ thermo/nociceptors (reddish colors) in WT, *Neurog2^1/1^* and *Neurog1^2/2^*embryos at E18.5. Numbers inside the graphics indicate the percentage of each neuronal subtypes. Blue and red numbers outside the graphics indicate the relative proportions of mechano/proprioceptors *vs.* thermo/nociceptors, respectively.

Between E9.5 and E11, expression of *Neurog1* and *Neurog2* largely overlap. Yet, *Neurog2* is uniquely induced in migrating somatosensory progenitors (Ma et al., 1999), suggesting that proper activity of the *Neurog2* locus in this cell population is critical during this process. To test this hypothesis, we took advantage of the *2^1neo1/1neo^* transgenic model in which the *Neurog2* locus is correctly activated at E9.5 but prematurely turned off by E10.5, while the activity of the *Neurog1* locus in post-migratory progenitors remains largely unaffected (Figure S3E). In these transgenic embryos, we found a significant increase in the number of *Mitf*+ melanoblasts compared to WT, strikingly similar to that observed in *Neurog2^-/-^* KO (Figures 2D,E,I,N). An identical phenotype was also observed in *2^1neo1/1neo^*;*Neurog1^-/-^*compound transgenic models (Figures 2J,N), further illustrating that the *Neurog1* locus activity is not required. These results thus establish that precise spatiotemporal regulation of the *Neurog2* locus activity in migrating progenitors is essential for committing them to the somatosensory lineage and preventing their differentiation into melanoblasts.

### *Neurog1* is slightly less effective than *Neurog2* in inhibiting a melanocyte identity *in vivo* and *in vitro*

During the course of this study, we uncovered a dose-dependent requirement for *Neurog2* in repressing the melanocyte identity. Indeed, in E11.5 *Neurog2^+/-^* heterozygous embryos, we detected a higher number of *Mitf*+ melanoblasts compared to WT, which nevertheless remained far below that of *Neurog2^-/-^*homozygous mutants (Figures 2D,E,K,N). This prompted us to also analyze *Neurog2^1/-^*hemizygous animals in which one copy of the endogenous *Neurog2-CDS* is replaced by the *Neurog1-CDS*, while the other copy is deleted. In this model, we found a significant increase in the number of melanoblasts, not only compared to WT but also compared to *Neurog2^+/-^* heterozygous embryos (Figures 2D,E,K,L,N). In contrast, in *Neurog2^1/+^* heterozygous KI animals in which one copy of the *Neurog2-CDS* and one copy of the *Neurog1-*CDS are expressed from the endogenous *Neurog2* loci, no supernumerary melanoblasts were detected (Figures 2M,N). These findings illustrate that, beyond its spatiotemporal regulation, the level of activity of the *Neurog2* locus is also a key parameter. Importantly, they also provide evidence that *Neurog1* is slightly less efficient than *Neurog2* in repressing the melanocyte fate, revealing subtle functional differences between the two TFs.

To gain further insights into this observation, we conducted in vitro analyses using the B16F10 melanoma cell line. We first transfected varying concentrations of *Neurog1* or *Neurog2* expression vectors and quantified the amount of *MitF* mRNA by RT-qPCR 24h later. We found that at a concentration of 500ng/µl, *Neurog1* and *Neurog2* were equally able to significantly reduce *MitF* expression, while at a lower concentration of 200ng/µl, only *Neurog2* was efficient in triggering such down-regulation (Figure 2O). To refine this analysis, we co-transfected a *gfp*-vector with either the *Neurog1*-vector or the *Neurog2*-vector at a concentration of 200ng/µl and quantified the proportion of GPF+ cells that were Mitf+ or Mitf-negative (-). At this concentration, we found that *Neurog2*, but not *Neurog1*, was able to significantly decrease the proportion of GFP+/Mitf+ cells compared to the control condition (Figure 2P-S). These data show that both *Neurog1* and *Neurog2* can down-regulate *Mitf in vitro*. They also confirm that *Neurog1* is not as efficient as *Neurog2* during this process.

Although Neurog1 and Neurog2 are considered as close paralogous TFs, their bHLH domains are not identical, and their N- and C-terminus domains are highly divergent (Figure S2A; Hand et al., 2005). This prompted us to investigate whether the functional discrepancies between the two TFs stem from divergences in their bHLH domains and/or in other regions of the proteins. To test this, we designed chimeric versions of Neurog1 and Neurog2 with swapped bHLH domains (Figure S2B), and overexpressed them at a concentration of 200ng/µl in B16F10 cells. Strikingly, we found that the Neurog1-bHLH2 chimera was as efficient as the native Neurog2 protein to decrease the proportion of GFP+Mitf+ cells, compared to the control or “native Neurog1” conditions (Figures 2P,Q,S,T). Nevertheless, the Neurog2-bHLH1 chimera had also a significant, albeit milder, effect (Figure 2P,Q,U). These results thus support that the slight functional differences between Neurog1 and Neurog2 in repressing the melanocyte identity are primarily due to divergences in their bHLH sequences, although contribution of other domains cannot be ruled out.

### Four copies of either *Neurog1-CDS* or *Neurog2-CDS* ensure proper neurogenesis and fate specification in the developing DRG

*Neurog1* and *Neurog2* act in a combinatorial manner to regulate the successive waves of DRG neurogenesis which generate specific neuron numbers and functional classes (Figure S1A; Ma et al., 1999, Ohayon et al., 2015; Venteo et al., 2019; Vermeiren et al., 2020). We thus investigated whether exchanging the *Neurog1*-CDS and the *Neurog2-*CDS in *Neurog1^2/2^* and *Neurog2^1/1^* KI models might affect the number and identities of the neurons generated or, conversely, ensure proper DRG development and rescue the defects of *Neurog1^-/-^*and *Neurog2^-/-^* KO models. First, we monitored neurogenesis through time course analyses of *NeuroD* and *SCG10* expression between E10.5 and E13.5, which identify differentiating precursors and newly born post-mitotic neurons, respectively. On one hand, in *Neurog2^1/1^* KI embryos we found that neurogenesis was initiated on time, as evidenced by normal numbers of *NeuroD*+ and *SCG10*+ precursors at E10.5, in contrast to the delay observed in the *Neurog2^-/-^*KO (Figures 3A-C; Ma et al., 1999; Venteo et al., 2019). Furthermore, the rate of neurogenesis over time in the KI was strikingly similar to WT, leading to the production of accurate numbers of neurons at E13.5, which contrasted with the 30% deficit reported in *Neurog2^-/-^* KO (Figures 3A,B; Venteo et al., 2019). On the other hand, in *Neurog1^2/2^* KI embryos, neurogenesis also proceeded normally over time and at E13.5 their DRG contained appropriate neuron numbers (Figures 3D,E). This differed markedly from the premature arrest of neurogenesis at E11.5 and the resulting 80% neuronal deficit observed in the *Neurog1^-/-^* KO (Figures 3D,E; Ma et al., 1999; Ohayon et al., 2015). Notably, at E12.5 when no *NeuroD*+ differentiating precursors and fewer *SCG10*+ neurons were detected in *Neurog1^-/-^*KO, both populations were clearly present in *Neurog1^2/2^* KI embryos, as in WT (Figure 3F). These data thus show that four copies of either the *Neurog1*-CDS or *Neurog2*-CDS faithfully control the successive waves of neurogenesis and rescue the defects caused by *Neurog1* or *Neurog2* loss of function. They also indicate that *Neurog1* and *Neurog2* do not directly and differentially regulate the number of neurons generated from the different NC-derived somatosensory progenitor populations (Frank and Sanes, 1991; Marmigere et al., 2007; Ohayon et al., 2015).

Next, we assessed the identities of the somatosensory neurons generated in the *Neurog1^2/2^* and *Neurog2^1/1^* KI models. We primarily focused on E13.5, at the end of DRG neurogenesis, and E18.5, at the end of embryogenesis, as *Neurog1^-/-^* and *Neurog2^-/-^* KO models die around birth. At E13.5, we classically used the tyrosine kinase receptor TrkA to identify thermo/nociceptive precursors, and TrkB and TrkC to identify mechano/proprioceptive precursors. At E18.5, although neuronal diversification and maturation are still ongoing, the expression of several molecular markers already identify specific subtypes within each functional class. Those include *TrkB*, *TrkC* and *MafA* for mechanoceptors, *TrkC* and Parvalbumin (PV) for proprioceptors, and *MrgprD*, *TrpV1*, *TrpM8* or *Cgrp* for Thermo/Nociceptors (Lallemend and Ernfors, 2012; Usoskin et al., 2015; Sharma et al., 2020; Cranfill & Luo, 2021). Analysis of this set of markers in *Neurog1^2/2^* and *Neurog2^1/1^*KI animals revealed that all somatosensory neuron subtypes were present in normal numbers compared to controls at both E13.5 (Figure 3G) and E18.5 (Figure 3H). Notably, while *Neurog1* is necessary to generate most thermo/nociceptors (Figures S1B; see also Figures 4D,F,G; Ma et al., 1999; Ohayon et al., 2015), its sole expression in *Neurog2^1/1^* KI embryos was not sufficient to trigger the overproduction of this neuronal class. Accordingly, in both KI models the relative distribution of each neuron subtype as well as the overall proportions of Mechano/Proprioceptors *versus* Thermo/Nociceptors were fully comparable to WT (Figure 3I). These results thus show that four copies of either the *Neurog1* or *Neurog2* CDS can efficiently support the differentiation of all DRG neuron subtypes, establishing that both paralogous TFs do not preferentially specify distinct neuronal identities.

**Figure 4.**
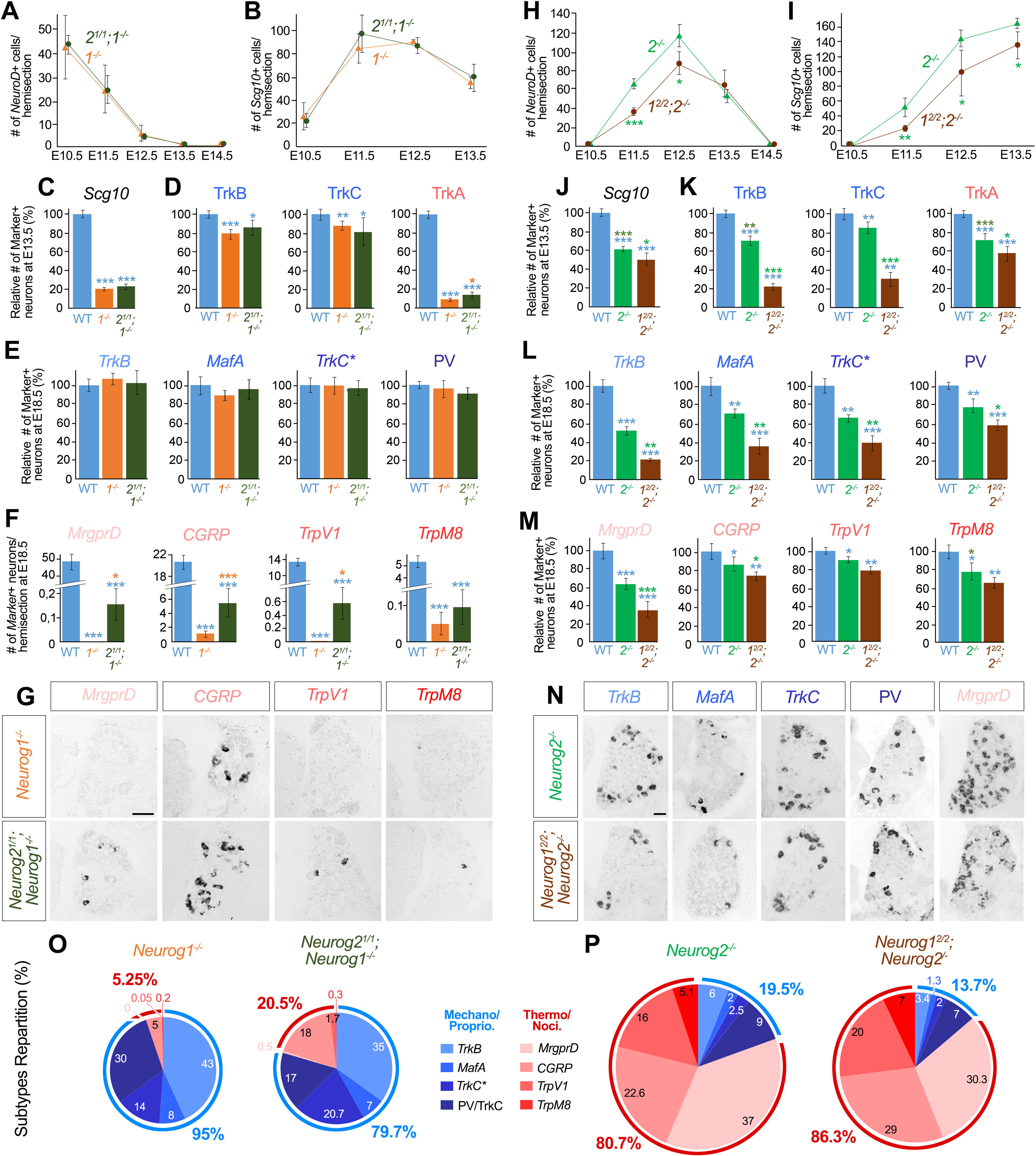
Specific combinations of KI and KO alleles uncover subtle functional differences between Neurog1 and Neurog2 during DRG neuron differentiation. **A,B.** Quantitative time-course analysis of *NeuroD*+ differentiating precursors between E10.5 and E14.5 (A) and *Scg10*+ post-mitotic neurons between E10.5 and 13.5 (B) in *Neurog1^-/-^* KO (*1^-/-^*; orange curves) and *Neurog2^1/1^*;*Neurog1^-/-^* KI;KO models (*2^1/1^*;*1^-/-^* dark green curves) showing no significant difference between the two genotypes over time. Data are represented as means ± SEM. **C,D.** Relative quantification of *Scg10*+ (C) and TrkB+, TrkC+ and TrkA+ (D) neuronal precursors in the DRG of WT (blue columns), *Neurog1^-/-^*KO (orange columns) and *Neurog2^1/1^*;*Neurog1^-/-^*KI;KO (*2^1/1^*;*1^-/-^* dark green columns) embryos at E13.5. Data are represented as means ± SEM. *p < 0.05; ***p < 0.001. Blue asterisks indicate significant differences with WT. Orange asterisks indicate significant differences with *Neurog1^-/-^* KO. **E.** Relative quantification of the *TrkB*+, *MafA*+, *TrkC*+/PV- (*TrkC\**) and PV+ mechano/proprioceptive subtypes in WT (blue columns), *Neurog1^-/-^*KO (orange columns) and *Neurog2^1/1^*;*Neurog1^-/-^*KI;KO (*2^1/1^*;*1^-/-^* dark green columns) embryos at E18.5. Data are represented as means ± SEM. **F.** Quantification of the numbers of *MrgprD*+, *Cgrp*+, *TrpV1*+ and *TrpM8*+ thermo/nociceptive subtypes, reported by hemi-section, in WT (blue columns), *Neurog1^-/-^* KO (orange columns) and *Neurog2^1/1^*;*Neurog1^-/-^* KI;KO (*2^1/1^*;*1^-/-^* dark green columns) embryos at E18.5. Data are represented as means ± SEM. *p < 0.05; ***p < 0.001. Blue asterisks indicate significant differences with WT. Orange asterisks indicate significant differences with *Neurog1^-/-^*KO. **G.** Illustrative images of *in situ* hybridization for *MrgprD*, *Cgrp*, *TrpV1* and *TrpM8* on transverse DRG sections of *Neurog1^-/-^*KO and *Neurog2^1/1^*;*Neurog1^-/-^* KI;KO embryos at E18.5. Scale bar, 50µm. **H,I.** Quantitative time-course analysis of *NeuroD*+ differentiating precursors between E10.5 and E14.5 (A) and of *Scg10*+ post-mitotic neurons between E10.5 and 13.5 (B) in *Neurog2^-/-^*KO (*2^-/-^*; green curves) and *Neurog1^2/2^*;*Neurog2^-/-^*KI;KO embryos (*1^2/2^*;*2^-/-^* dark brown curves) revealing exacerbated neurogenesis defects at E11.5 and E12.5 in the KI;KO model. Data are represented as means ± SEM. *p < 0.05; ***p < 0.001. Green asterisks indicate significant differences with *Neurog2^-/-^* KO. **J,K.** Relative quantification of *Scg10*+ (J) and TrkB+, TrkC+ and TrkA+ (K) neuronal precursors in the DRG of WT (blue columns), *Neurog2^-/-^*KO (green columns) and *Neurog1^2/2^*;*Neurog2^-/-^*KI;KO (*1^2/2^*;*2^-/-^* dark brown columns) embryos at E13.5. Data are represented as means ± SEM. *p < 0.05; **p < 0.01; ***p < 0.001. Blue asterisks indicate significant differences with WT. Green asterisks indicate significant differences with *Neurog2^-/-^* KO. **L,M**. Relative quantification of *TrkB*+, *MafA*+, *TrkC*+/PV- (*TrkC\**) and PV+ mechano/proprioceptive subtypes (L) and *MrgprD*+, *Cgrp*+, *TrpV1*+ and *TrpM8*+ thermo/nociceptive (M) subtypes in WT (blue columns), *Neurog2^-/-^* KO (*2^-/-^*; green columns) and *Neurog1^2/2^*;*Neurog2^-/-^*KI;KO (*1^2/2^*;*2^-/-^*dark brown columns) embryos at E18.5. Data are represented as means ± SEM. *p < 0.05; **p < 0.01; ***p < 0.001. Blue asterisks indicate significant differences with WT. Green asterisks indicate significant differences with *Neurog2^-/-^* KO. **N.** Illustrative images of *in situ* hybridization for *TrkB*, *MafA*, *TrkC* and *MrgprD*, or immunostaining for PV, on transverse DRG sections of *Neurog2^-/-^* KO and *Neurog1^2/2^*;*Neurog2^-/-^* KI;KO embryos at E18.5 showing reduced numbers of positive neurons for all these markers in the KI;KO model compared to KO. Scale bar, 50µm. **O,P.** Graphic summaries showing the distribution of the *TrkB*+, *MafA*+, *TrkC*+/PV- (*TrkC\**) and PV+ mechano/proprioceptive subtypes (blue colors) and the *MrgprD*+, *CGRP*+, *TrpV1*+ and *TpM8*+ thermo/nociceptive subtypes (reddish colors) in the DRG of *Neurog1^-/-^*KO and *Neurog2^1/1^*;*Neurog1^-/-^* KI;KO embryos (O), and of *Neurog2^-/-^* KO and *Neurog1^2/2^*;*Neurog2^-/-^*KI;KO embryos (P) at E18.5. Numbers inside the graphics indicate the percentage of each neuronal subtypes. Blue and red numbers outside the graphics indicate the relative proportions of mechano/proprioceptors *vs.* thermo/nociceptors in each model, respectively. Note that the neuronal distribution between the KO and the corresponding KI;KO models are changed.

Taken altogether, these data show that in the developing DRG, *Neurog1* and *Neurog2* are essentially functionally interchangeable during the processes of neurogenesis and fate specification, further illustrating that their apparent functional specificities primarily stem from differences in their spatio-temporal expression profiles.

### Combinations of KI and KO alleles reveal *s*ubtle functional differences between Neurog1 and Neurog2 in DRG neuron development

To complement this study, we analyzed *Neurog2^1/1^*;*Neurog1^-/-^*and *Neurog1^2/2^*;*Neurog2^-/-^* KI;KO in comparison to *Neurog1^-/-^* KO and *Neurog2^-/-^* KO embryos, respectively. These models all express only two copies of the *Neurog1-CDS* or the *Neurog2*-CDS, either from the endogenous loci in the KO (Figures 1P,S) or from the counterpart loci in the KI;KO (Figures 1R,U). In the light of the melanocyte data, we aimed to uncover potential functional divergences between the two TFs that might be masked in single-KI models, considering the transient overlapping activities of the *Neurog1* and *Neurog2* gene loci between E9.5 and E11.5 (Figures 1M-O and S1A).

On one hand, comparative time course analysis of *NeuroD* and *Scg10* expression between *Neurog2^1/1^*;*Neurog1^-/-^* KI;KO and *Neurog1^-/-^* KO embryos showed that the timing and rate of neurogenesis over time were similar in both models, with a normal onset at E9.5 but a premature end at E11.5 (Figures 4A,B; Ma et al.1999; Ohayon et al., 2015). As a result, at E13.5, the overall number of *Scg10*+ neurons was equally reduced by approximately 80% in both models compared to controls (Figures 4B,C and S4A). Analyses of neuron identities at E13.5 revealed a relatively mild reduction of the numbers of TrkB+ and TrkC+ mechano/proprioceptive precursors in both KI;KO and KO embryos compared to WT (Figures 4D and S4A), which was, however, no longer evident at E18.5 (Figures 4E and S4B). In contrast, the number of TrkA+ thermo/nociceptive precursors at E13.5 was drastically reduced by 93% in the *Neurog1^-/-^* KO and by 86% in the *Neurog2^1/1^*;*Neurog1^-/-^* KI;KO embryos, compared to WT (Figures 4D and S4A). Strikingly, this depletion was significantly less severe in the KI;KO than in the single-KO (Figure 4D), a result also observed at E18.5. At this stage, in *Neurog1^-/-^* single-KO embryos (n=5), virtually no *MrgprD*+ (0/5) or *TrpV1*+ (0/5), and only few, if any, *TrpM8*+ (1/5) neurons were present (Figures 4F,G). The only remaining thermo/nociceptors in the KO were *Cgrp*+ (Figures 4F,G) and likely derived from the first neurogenic wave (Figure S1B; Bachy et al., 2011; Lallemend and Ernfors, 2012). In contrast, in *Neurog2^1/1^*;*Neurog1^-/-^*KI;KO embryos (n=6), we systematically detected few *MrgprD*+ (5/6), *TrpV1*+ (6/6) and *TrpM8*+ (5/6) neurons, as well as a significant increase of the *Cgrp*+ population (6/6) compared to *Neurog1^-/-^*KO (Figure 4F,G). Consequently, the relative distribution of neuron subtypes and the proportions of mechano/proprioceptors versus thermo/nociceptors differed between the KI;KO and KO models (Figure 4O). This phenotypic difference remains relatively marginal, affecting only 3% of the entire DRG population, and the underlying mechanism remains unclear. It may reflect the ability of *Neurog1* to impose a thermo/nociceptive identity on a small subset of progenitors that would normally generate mechano/proprioceptors under the control of *Neurog2*, although no difference in the latter population was observed between the KI;KO and KO models (see Figures 4D,E). Alternatively, it may result from a slight temporal shift of neurogenesis in the KI;KO compared to the KO which may influence neuron subtype specification (see below). In any case, these data highlight subtle functional differences between *Neurog1* and *Neurog2* in regulating DRG neuron differentiation.

On the other hand, comparative studies of *Neurog1^2/2^*;*Neurog2^-/-^*KI;KO and *Neurog2^-/-^* KO models led to similar observations. Time course analysis of *NeuroD* and *Scg10* expression showed that onset of neurogenesis was delayed by 24 hours in both models, beginning after E10.5 instead of E9.5 (Figures 4H,I; Ma et al., 1999; Venteo et al., 2019). Once initiated, however, we found that neurogenesis proceeded at a lower rate in the KI;KO compared to KO embryos, notably between E11.5 and E12.5 (Figure 4H). Consequently, *Scg10*+ post-mitotic precursors accumulated in fewer numbers in *Neurog1^2/2^*;*Neurog2^-/-^* KI;KO embryos over time (Figure 4I), leading to an overall neuronal depletion at E13.5 of approximately 20% compared to *Neurog2^-/-^* KO, and 50% compared to WT (Figures 4J and S4C). In addition, analysis of the neuronal identities at E13.5 showed a decrease of all neuronal classes in the KI;KO compared to the KO, albeit more pronounced for the TrkB+ and TrkC+ mechano/proprioceptive precursors than the TrkA+ thermo/nociceptive precursors (Figures 4K and S4C), a result also confirmed at E18.5. At this stage, the depletion of all mechano/proprioceptive subtypes reported in the *Neurog2^-/-^* KO, was exacerbated in the *Neurog1^2/2^*;*Neurog2^-/-^* KI;KO embryos, although we noticed that the *TrkB*+, *MafA*+, *TrkC*+ mechanoceptors appeared slightly more affected than the PV+ proprioceptors (Figures 4L,N). In the same line, if all thermo/nociceptive subtypes showed at least a tendency to decrease in the KI;KO compared to the KO, the *MrgprD*+ population appeared particularly more impacted (Figures 4M,N and S4D). As a consequence, the distribution of neuron subtypes, as well as the relative proportions of mechano/proprioceptors versus thermo/nociceptors, were altered in the *Neurog1^2/2^*;*Neurog2^-/-^*KI;KO compared to the *Neurog2^-/-^* KO, with a stronger impact on the mechano/proprioceptive class (Figure 4P).

Taken together, these findings uncover subtle functional discrepancies between the Neurog1 and Neurog2 proteins in regulating DRG neurogenesis which are, however, masked by dose- compensation effects, as no neuronal defects were detected in *Neurog2^1/1^* and *Neurog1^2/2^* single-KI models (see Figure 3).

### Mechano/Proprioceptive and Thermo/nociceptive subtypes are preferentially born during distinct developmental time-windows

The fact that *Neurog1* and *Neurog2* are essentially functionally interchangeable in controlling fate specification in the forming DRG notably indicates that *Neurog1* is not uniquely involved in specifying the thermo/nociceptive identity. Therefore, the specific loss of the vast majority of this neuronal class in *Neurog1^-/-^* KO, which correlates with impaired neurogenesis from E11.5 (Figures 4 and S1B), implies that mechano/proprioceptors and thermo/nociceptors are primarily born during distinct developmental periods. Previous birth-dating studies have indeed reported successive peaks of birth for both neuronal classes (Lawson and Biscoe, 1979; Kitao et al., 1996). However, this view has been challenged by a more recent study suggesting that many thermo/nociceptors are already generated at E10.5, concurrently with mechano/proprioceptors (Landy et al., 2021). These contradictory outcomes prompted us to conduct a new series of birth-dating experiments. To do so, we injected BrdU into pregnant mice at E8.5, E9.5, E10.5, E11.5, E12.5, E13.5 or E14.5, harvested the embryos at E18.5 (for consistency with analyses on KO and KI models), and performed double-labeling experiments using BrdU with either *TrkB*, *TrkC*, *MafA*, PV, *MrgprD*, *Cgrp*, *TrpV1* or *TrpM8* as subtypes markers, or with *Scg10* as a pan-neuronal marker (Figure S5A). As a prerequisite, we ensured that all “Marker X+” (X+) populations analyzed were properly represented in our experimental paradigm, by verifying that the distribution of each X+/BrdU+ subtype relative to the total number of *Scg10*+/BrdU+ neurons across all experiments was comparable to the normal distribution of each X+ subtype relative to the total number of *Scg10*+ DRG neurons in E18.5 WT embryos (Figure S5B). We then evaluated the global temporal pattern of neuron production by determining the proportion of *Scg10+*BrdU+ neurons after each BrdU injection, relative to the total number of *Scg10*+BrdU+ neurons counted across all experiments. As expected, we found that DRG neuron differentiation began at E9.5, peaked between E11.5 and E12.5, and ended after E13.5 (Figure 5A). We also confirmed that at E9.5- E11.5 and E11.5-E13.5, encompassing the two main neurogenic phases, approximately 25% and 75% of neurons were generated, respectively (Figure 5A; summarized in 5F). Next, we assessed the birth of the different mechano/proprioceptive and thermo/nociceptive subtypes over time. Given the variable size of each population, we first determined the raw number of each X+BrdU+ subtypeat E18.5 after each BrdU injection (Figures 5B). Using this specific combination of molecular markers, we observed that although most neuronal subtypes were continuously born between E9.5 and E13.5, mechano/proprioceptors were preferentially generated at E9.5 and E10.5, while thermo/nociceptors were predominantly born between E11.5 and E13.5 (Figure 5B). Accordingly, by separately grouping mechano/proprioceptors and thermo/nociceptors and assessing their relative proportions at each time point, we found that 93% and 76% of the neurons generated at E9.5 and E10.5, respectively, were mechano/proprioceptors, whereas approximately 95% of the neurons produced at each time point from E11.5 were thermo/nociceptors (Figure 5C; summarized in 5F). Next, we assessed the birth rate of each X+ subtype over time by analyzing the percentage of X+BrdU+ neurons after each BrdU injection, relative to the total number of X+BrdU+ neurons across all experiments. Thereby, we found that all subtypes within a given class exhibited a similar peak of birth: between E10.5 and E11.5 for mechano/proprioceptors and between E11.5 and E12.5 for thermo/nociceptors (Figure 5D). Furthermore, by separately grouping each neuronal class, we determined that about 85% of the mechano/proprioceptors were born between E9.5 and E11.5, while 95% of the thermo/nociceptors were born between E11.5 and E13.5 (Figure 5E; summarized in 5F).

**Figure 5.**
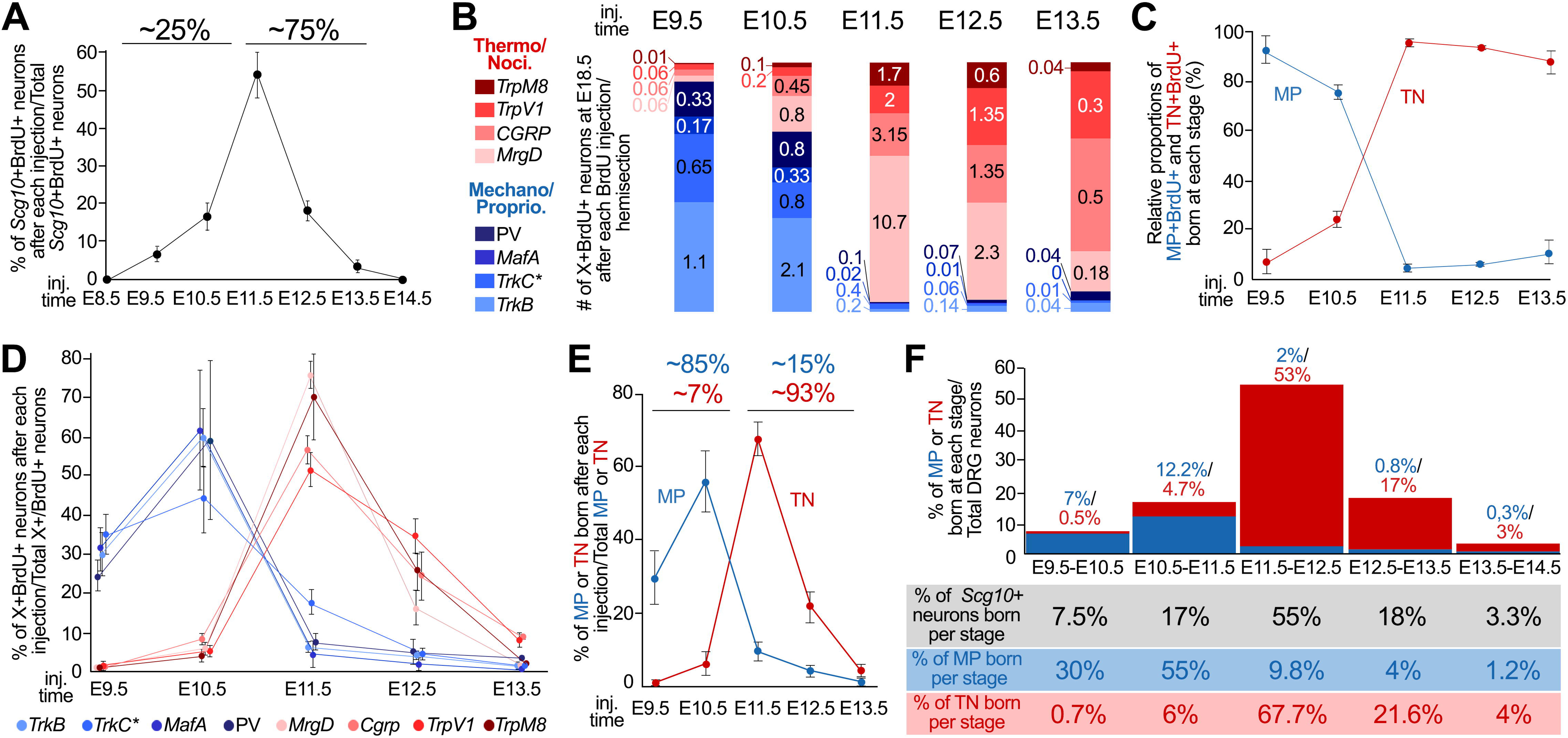
Mechano/proprioceptive and thermo/nociceptive subtypes are preferentially born during successive developmental periods. **A.** Quantification of the proportion of *Scg10*+ neurons born at each stage between E8.5 and E14.5, showing that: (i) DRG neurogenesis proceeds between E9.5 and E13.5; (ii) the peak of DRG neuron production occurs between E11.5 and E12.5; and (iii) during the two main neurogenic waves of the DRG, at E9.5-E11.5 and E11.5-E13.5, approximately 25% and 75% of the DRG neurons are produced, respectively. Data are represented as means ± SEM. **B.** Numbers of *TrkB*+/BrdU+, *MafA*+/BrdU+, *TrkC*+/BrdU+, PV+/BrdU+, *MrgprD*+/BrdU+, *Cgrp*+/BrdU+, *TrpV1*+/BrdU+ and *TrpM8*+/BrdU+ (globally referred to as X+/BrdU+) neurons at E18.5, reported by hemisection, after each BrdU injection between at E9.5 and E13.5, as indicated. This shows that although all subtypes are produced continuously over time, all mechano/proprioceptors (blue bars) and all thermo/nociceptors (reddish bars) are predominantly produced during distinct periods. **C.** Relative proportions of mechano/proprioceptors (MP, blue curve) *vs.* thermo/nociceptors (TN, red curve) generated at each stage between E9.5 and E13.5. Data are represented as means ± SEM. **D.** Quantitative analysis of the birth rate of each X+ subtype over time, determined by calculating the proportions of X+BrdU+ neurons at E18.5 after each BrdU injection between E9.5 and E13.5, relative to the total number of X+BrdU+ neurons across all experiments. This shows that all analyzed mechano/proprioceptive subtypes (blue curves) and thermo/nociceptive subtypes (reddish curves) exhibit successive peaks of birth: E10.5-E11.5 for the former and E11.5-E12.5 for the latter. Data are represented as means ± SEM. **E.** Proportions (i) of mechano/proprioceptors generated at each stage between E9.5 and E13.5, relative to the whole population of mechano/proprioceptors (blue curve), and (ii) of thermo/nociceptors generated at each stage relative to the whole population of thermo/nociceptors. This further that each class of neurons exhibit successive birth peaks and establishes that approximately 85% of mechano/proprioceptors are born before E11.5, whereas 93% of thermo/nociceptors are born after E11.5. Data are represented as means ± SEM. **F.** Summary of the birth dating data. The graph illustrates the proportions of mechano/proprioceptors (MP, blue columns and numbers) and thermo/nociceptors (TN, red columns and numbers) generated at each stage between E9.5 and E13.5, relative to the total numbers of *Scg10*+ DRG neurons at E18.5. The table below recapitulates the percentages of *Scg10*+ neurons (see A), of mechano/proprioceptors (MP; see E) and of thermo/nociceptors (TN; see E) born at each stage between E9.5 and E13.5 relative to the whole *Scg10*+ population, MP population and TN population, respectively.

Altogether, these data establish that the majority of the mechano/proprioceptive and thermo/nociceptive subtypes are preferentially born during successive developmental periods. It is of note that this analysis excluded some subtypes, notably the population of C-fiber low threshold mechanoceptors known to emerge from the thermo/nociceptive lineage (Lou et al., 2013; Vermeiren et al. 2021), and are likely born later than the “classical” mechanoceptors assessed here (see Landy et al., 2021). However, despite this drawback, our findings are fully consistent with previous studies supporting that the successive neurogenic waves in the DRG generate 20% and 80% of the DRG neurons, mainly mechano/proprioceptors and thermo/nociceptors, respectively (Figure S1A; Lawson and Biscoe, 1979; Kitao et al., 1996; Ma et al., 1999; Marmigere et al., 2007; Lallemend and Ernfors; 2012; Ohayon et al., 2015; Vermeiren et al., 2021).

### Defined alterations of neurogenesis predictably alter DRG content, highlighting the critical influence of differentiation timing in biasing neuronal fates

Birth-dating and functional studies notably on *Neurog1^-/-^*KO, *Neurog2^-/-^* KO and *Neurog1^2/2^;Neurog2^-/-^* KI;KO models suggest a strong correlation between the timing and rate of differentiation and the identities and numbers of somatosensory neurons generated. This prompted us to assess whether defined alterations of neurogenesis can have predictable consequences on the DRG content. To address this issue, we first retrospectively analyzed *Neurog1^-/-^* KO*, Neurog2^-/-^*KO and *Neurog1^1/1^;Neurog2^-/-^* KI;KO which exhibit distinct and specific neurogenic defects (see Figure 4). In these models, we determined the ratio of neurogenesis at each stage between E9.5 and 13.5 relative to WT (set to 1), by calculating the relative number of neurons produced over time compared to control embryos (Figure 6A). Based on these ratios and the birth-dating quantifications summarized in the table of Figure 5F, we estimated for each model the proportions of overall *Scg10*+ neurons, of mechano/proprioceptors (MP) and of thermo/nociceptors (TN) generated at each stage and, in turn, the relative numbers of each neuronal population predicted to ultimately reside in their DRG compared to controls (Figure 6B-D). Particularly for the *Neurog2^-/-^* KO and *Neurog1^1/1^;Neurog2^-/-^*KI;KO models, experimental data collected at E18.5 strikingly matched the predicted decreases in the numbers of overall *Scg10*+ neurons (29.9% *vs.* 33.5% and 51.8% *vs.* 54.5%, respectively), of thermo/nociceptors (37.3% *vs.* 50.3%, and 70.5% *vs.* 72.4%, respectively) and of mechano/proprioceptors (24.8% *vs.* 29% and 47.8% *vs.* 49.8%, respectively) (Figures 6B-C’). In *Neurog1^-/-^* KO, the drastic reduction in the numbers of overall *Scg10*+ neurons and thermo/nociceptors at E18.5 also closely aligned with the predictions (76% vs. 77.1% and 98% vs. 92%, respectively) (Figure 6D,D’). Intriguingly, however, the mechano/proprioceptive class, expected to be reduced by 27% in these mutants, were largely unaffected (Figure 6D,D’), although they were significantly decreased by 17% at E13.5 (see Figure 4D; brackets in Figure 6D’). The reasons for this phenotypic differences between the two stages remain unclear, but they may reflect: (i) a normalization effect of the naturally occurring cell death, (ii) a compensatory mechanism possibly involving an identity switch of a small population of thermo/nociceptive precursors toward the mechano/proprioceptive fate, and/or (iii) more dynamic neurogenic defects than reported so far. Nonetheless, aside from this exception, experimental data from these transgenic models generally aligned well with the predictions, establishing a close link between timing and rate of neurogenesis and DRG content (Summarized in Figures 7A-D).

**Figure 6.**
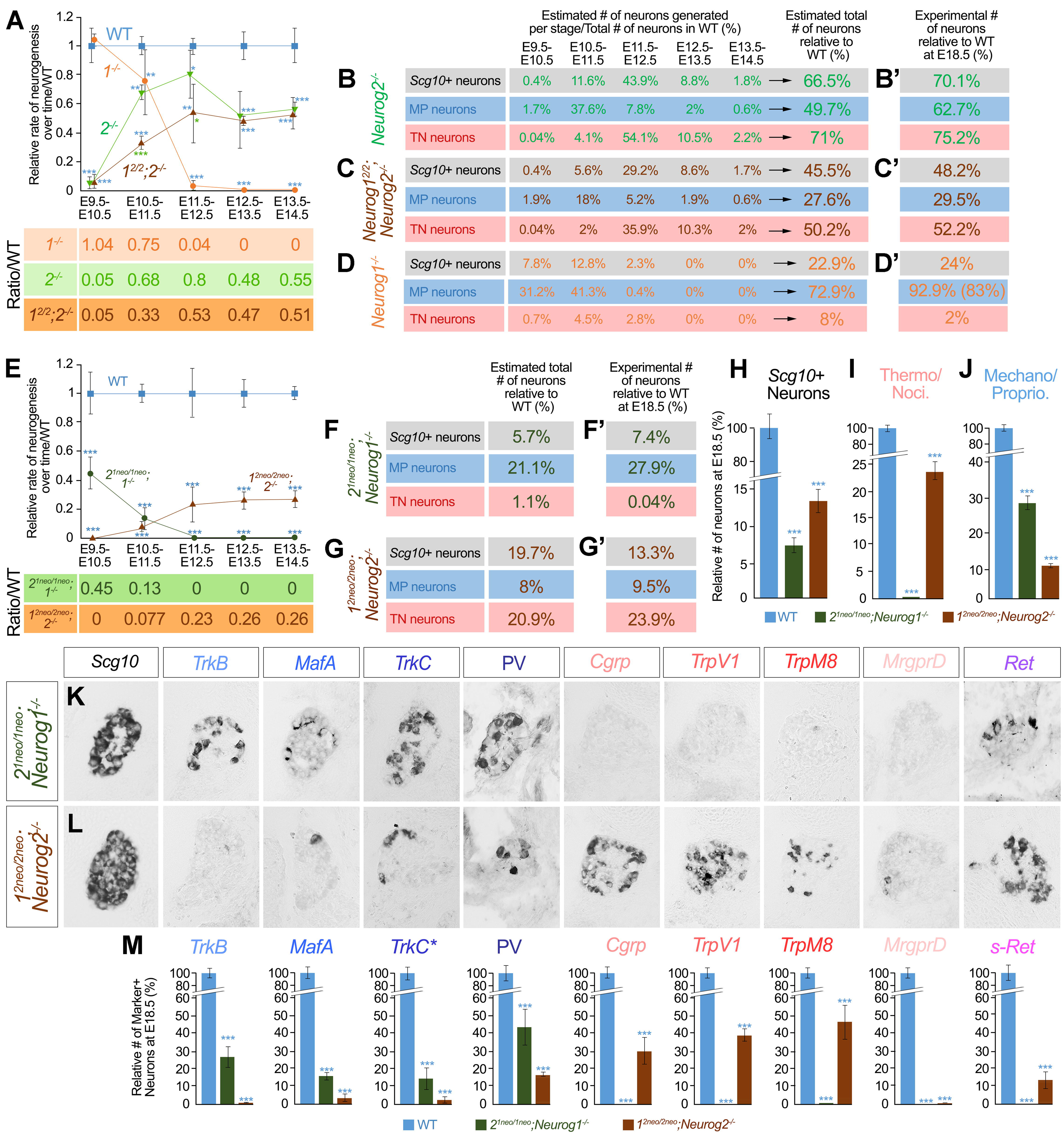
Defined alterations of the rate and timing of neurogenesis have predictable consequences on DRG content. **A.** Rate of neurogenesis at each stage between E9.5 and E13.5 in *Neurog1^-/-^* (*1^-/-^*; orange curve), *Neurog2^-/-^*(*2^-/-^*; green curve) and *Neurog1^2/2^;Neurog2^-/-^*(*1^2/2^;2^-/-^*; dark brown curve) models, relative to WT (set at 1; blue curve). Data are represented as means ± SEM. *p < 0.05; **p < 0.01; ***p < 0.001. Blue asterisks indicate significant differences with WT. Green asterisks indicate significant differences between *Neurog1^2/2^;Neurog2^-/-^*KI;KO and *Neurog2^-/-^* KO. For each model, the mean ratios of neurogenesis over time are indicated below. **B-D.** Proportions of *Scg10*+ neurons, mechano/proprioceptors (MP neurons), or thermo/nociceptors (TN neurons), predicted to be generated at each stage between E9.5 and E13.5 in *Neurog2^-/-^* (*2^-/-^*) (B), *Neurog1^2/2^;Neurog2^-/-^*(*1^2/2^;2^-/-^*) (C) and *Neurog1^-/-^* (*1^-/-^*) (D) models. This was calculated based on the neurogenic ratios determined in Figure 6A and on the birth-dating data shown in the table of Figure 5F. Adding the predicted numbers from all stages provided an estimation of the overall numbers of *Scg10*+ neurons, mechano/proprioceptors, and thermo/nociceptors in each model, relative to WT (set at 100%). **B’-D’.** Experimentally determined numbers of *Scg10*+ neurons, mechano/proprioceptors (MP neurons) and thermo/nociceptors (TN neurons) in the DRG of *Neurog2^-/-^* (*2^-/-^*) (B’), *Neurog1^2/2^;Neurog2^-/-^* (*1^2/2^;2^-/-^*) (C’) and *Neurog1^-/-^* (*1^-/-^*) (D’) embryos at E18.5 relative to WT, revealing a striking match with the predictions. The only exception concerned the mechano/proprioceptive population in *Neurog1^-/-^*KO (D’), although this population was reduced at E13.5 (number in brackets in D’). **E.** Ratio of neurogenesis at each stage between E9.5 and E13.5 in *2^1neo/1neo^;Neurog1^-/-^*(*2^1neo/1neo^;1^-/-^*; dark green curve) and *1^2neo/2neo^;Neurog2^-/-^*(*1^2neo/2neo^;2^-/-^*: dark brown curve) models, relative to WT (set to 1; blue curve), showing opposite temporal shifts and reduced neurogenic rates over time in both models compared to WT. Data are represented as means ± SEM. ***p < 0.001. Blue asterisks indicate significant differences with WT. For each model, the mean ratios of neurogenesis over time are indicated below. **F,G.** Relative numbers of *Scg10*+ neurons, mechano/proprioceptors (MP neurons), and thermo/nociceptors (TN neurons), predicted to reside in the DRG of *2^1neo/1neo^;Neurog1^-/-^* (F) and *1^2neo/2neo^;Neurog2^-/-^* (G) transgenic models, compared to WT. See text for details. **F’,G’.** Experimentally determined numbers of *Scg10*+ neurons, mechano/proprioceptors (MP neurons), and thermo/nociceptors (TN neurons), in *2^1neo/1neo^;Neurog1^-/-^*(D) and *1^2neo/2neo^;Neurog2^-/-^* (E) embryos at E18.5, relative to WT. **H-J.** Relative numbers of *Scg10*+ neurons (H), thermos/nociceptors (I) and mechano/proprioceptors (J) in WT (set at 100%; blue columns) *2^1neo/1neo^;Neurog1^-/-^*(dark green columns) and *1^2neo/2neo^;Neurog2^-/-^* (dark brown columns) embryos at E18.5. Data are represented as means ± SEM. *** p<0.001. Blue asterisks indicate significant differences with WT. **K,L.** Illustrative images of *in situ* hybridization for *Scg10*, *TrkB*, *MafA*, *TrkC*, *Cgrp*, *TrpV1 TpM8*, *MrgprD* and *Ret*, and immunostaining for PV, on thoracic DRG transverse sections of *2^1neo/1neo^;Neurog1^-/-^* (G) and *1^2neo/2neo^;Neurog2^-/-^*(H) embryos at E18.5 showing opposite defects in both models. **M.** Relative numbers of *TrkB*+, *MafA*+, *TrkC+*/PV- (*TrkC\**), PV*+* mechano/proprioceptors, and *Cgrp*+, *TrpV1*+ and *TpM8*+, *MrgprD*+, as well as of small-diameter *Ret*+ (s-*Ret*) thermo/nociceptors, in WT (set to 100%; blue columns), *2^1neo/1neo^;Neurog1^-/-^*(dark green columns) and *1^2neo/2neo^;Neurog2^-/-^* (dark brown columns) embryos at E18.5. Note that quantification of the large-diameter *Ret*+ mechanoceptors which are also *MafA*+ is not represented. Data are represented as means ± SEM. ***p < 0.001. Blue asterisks indicate significant differences with WT.

**Figure 7.**
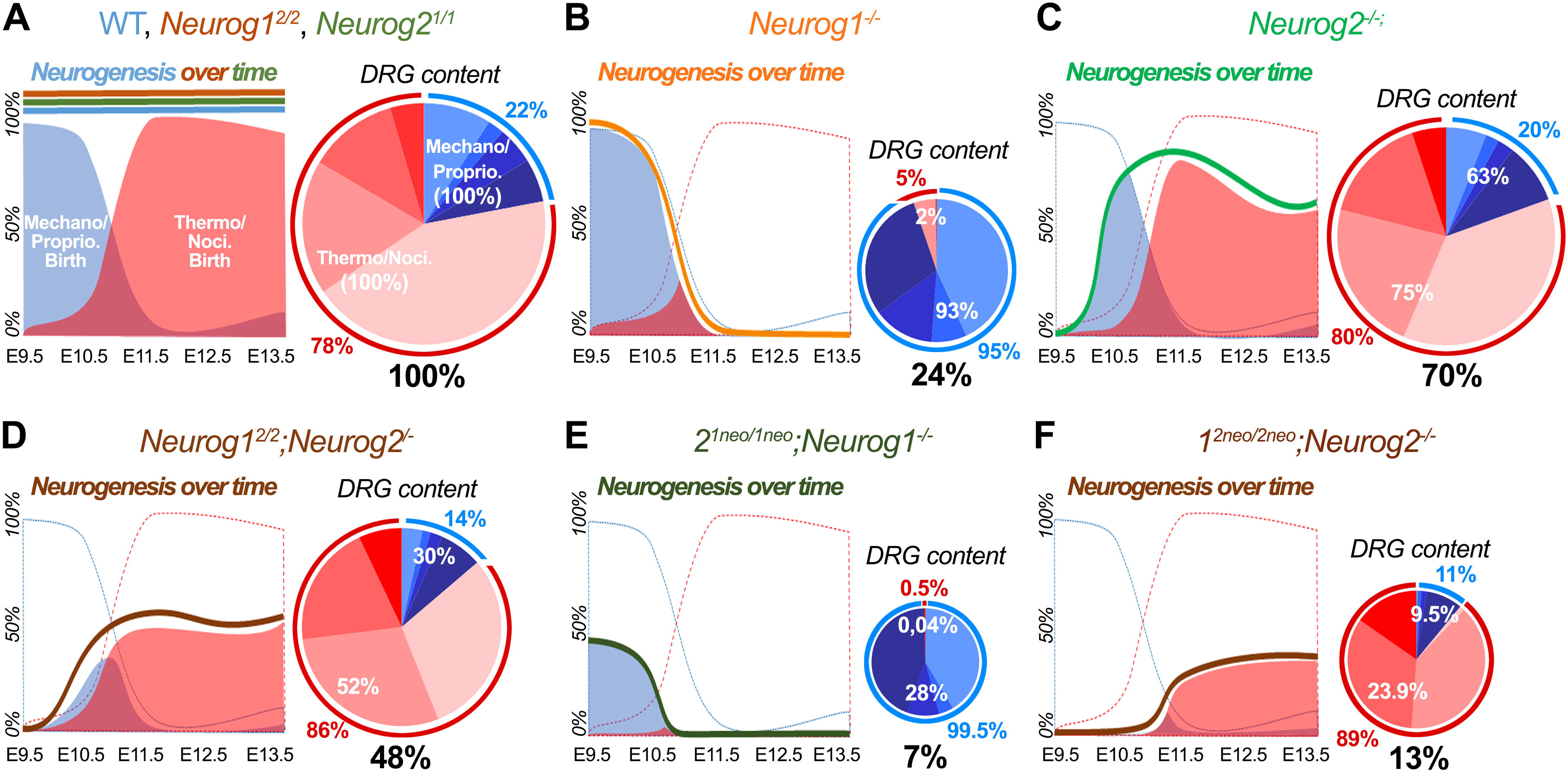
Summary of the phenotypes in transgenic mouse models linking timing and rate of neurogenesis, somatosensory neuron birth dates, and (iii) DRG content. **A-F.** Schematic summaries of the phenotypes of WT (A), *Neurog2^1/1^*single-KI (A), *Neurog1^2/2^* single-KI (A), *Neurog1^-/-^*single-KO (B), *Neurog2^-/-^* single-KO (C), *Neurog1^2/2^*;*Neurog2^-/-^* KI;KO (D), *2^1neo/1neo^;Neurog1^-/-^* (E) and *1^2neo/2neo^;Neurog2^-/-^* (F) models, illustrating a close link between timing and rate of neurogenesis over time, birth dates of mechano/proprioceptive and thermo/nociceptive neurons, and DRG content. For each model, the graph on the left shows the relative rate of neurogenesis between E9.5 and E13.5 (colored thick lines) compared to WT (set at 100%), in relation with the relative birth of mechano/proprioceptors (Mechano/Proprio.; blue-filled curve) *vs.* thermo/nociceptors (Thermo/Noci.; red-filled curve) over time. The pie chart on the right schematizes the DRG composition. The black percentage below each pie chart indicate the overall number of DRG neurons relative to WT (set at 100%). The white percentages inside each pie chart indicate the numbers of mechano/proprioceptors (Mechno/Proprio.) and thermo/nociceptors (Thermo/Noci.) relative to WT (set at 100%). The blue and red percentages outside each pie charts indicate the relative proportions of mechano/proprioceptors *vs.* thermo/nociceptors, respectively.

To complement this study, we took advantage of the *2^1neo/1neo^;Neurog1^-/-^*and *1^2neo/2neo^;Neurog2^-/-^* compound transgenic mouse models, in which the timing of *Neurog1/2* expression is altered in complementary ways, characterized by a premature down-regulation shortly after E10.5 in the former, and a delayed onset around E11.5 in the latter (see Figures 1Q,T). Accordingly, analysis of neurogenesis over time in these models, revealed opposite defects (Figure 6E). In *2^1neo/1neo^;Neurog1^-/-^*, neurogenesis began on time at E9.5, albeit at a lower rate compared to controls and *Neurog1^-/-^* KO, and ended prematurely shortly after E10.5 -approximately 48h and 24h earlier than in WT and *Neurog1^-/-^* KO, respectively (Figure 6E; compare with Figure 6A). Conversely, in *1^2neo/2neo^;Neurog2^-/-^*, neurogenesis started around E115, with a delay of approximately 48h and 24h compared to WT and *Neurog2^-/-^* KO, respectively. Moreover, from E11.5 to E13.5, neurogenesis proceeded at a lower rate compared to both WT and *Neurog2^-/-^*KO (Figure 6E; compare with Figure 6A). Estimations of the relative numbers of total DRG neurons, mechano/proprioceptors, and thermo/nociceptors generated in both models compared to WT, suggested that their DRG would be massively atrophied and predominantly composed of distinct neuronal classes (Figures 6F,G). On one hand, in the *2^1neo/1neo^;Neurog1^-/-^*model, we anticipated a decrease of 94.3% in the total number of neurons, 99% for the thermo/nociceptive class, and 78.9% for the mechano/proprioceptive class (Figure 6F). Experimental assessment of marker subtypes at E18.5 largely confirmed these predictions, showing a 92.6% reduction in the number of *Scg10*+ neurons (Figures 6F’,H,K), a nearly complete absence of thermo/nociceptors (Figures 6F’,I,K,M), and a 72.1% decrease in the number of mechano/proprioceptors (Figures 6F’,J,K,M). Therefore, temporally restricting the period of neurogenesis to E9.5-E10.5 particularly affected the thermo/nociceptive class, more drastically than in the *Neurog1^-/-^* KO, consistent with this population being primarily generated after E10.5 (Summarized in Figure 7E). Nevertheless, due to the reduced neurogenic rate at these stages compared to controls (Figures 6E), mechano/proprioceptive subtypes were also expectedly impacted (Figures 6F,F’,J,K,M and 7E). On the other hand, in the *1^2neo/2neo^;Neurog2^-/-^* model, we also observed a strong match between the experimentally determined, and the anticipated, decreases in the numbers of *Scg10*+ neurons (86.7% *vs.* 80.3%, respectively; Figures 6G,G’,H,L), thermo/nociceptors (76.1% *vs.* 77.2%, respectively; Figures 6G,G’,I,L,M) and mechano/proprioceptors (90.5% *vs.* 94.1%, respectively; Figures 6G,G’,J,L,M). Therefore, compared to the *Neurog2^-/-^* KO, further delaying the onset of neurogenesis, particularly affected the mechano/proprioceptive class, consistent with this population being mostly generated before E11.5 (Summarized in Figure 7F). However, again, due to the lower rate of neurogenesis in *1^2neo/2neo^;Neurog2^-/-^*embryos compared to *Neurog2^-/-^* KO, notably at E11.5- E12.5 (compare Figures 6A and 6E), the thermo/nociceptive class was also expectedly severely impacted (Figures 6 G,G’,I,L,M and 7F). It is of note that, considered individually, all neuron subtypes were not equally altered in the *1^2neo/2neo^;Neurog2^-/-^* embryos. Indeed, within the mechano/proprioceptive class, the PV+ proprioceptors appeared relatively less affected than the other subtypes (Figures 6L,M). In the same line, within the thermo/nociceptive class, while the *Cgrp*+, *TrpV1*+ and *TrpM8*+ populations were reduced but still present, very few, if any, *MrgprD*+ neurons were detected (Figures 6L,M). This population of small-diameter thermo/nociceptors also expresses the gene encoding the Ret receptor (Usoskin et al., 2015; Cranfill & Luo, 2021). Analysis of *Ret* expression in *1^2neo/2neo^;Neurog2^-/-^* embryos at E18.5, revealed the presence of significant numbers of small- sized *Ret*+ neurons (s-Ret, Figures 6L,M), suggesting that this neuronal population is produced but exhibits incomplete subtype-specific differentiation. Definite explanation for these observations remains elusive, but they may, at least in part, reflect developmental cooperation between the different precursor subtypes, as previously reported (Hadjab et al., 2013).

Taken altogether, these findings establish that opposite temporal shifts of neurogenesis -either delayed onsets in *1^2neo/2neo^;Neurog2^-/-^*or premature arrests in *2^1neo/1neo^;Neurog1^-/-^*-, have largely predictable and opposite impacts on DRG content, highlighting the critical importance of the timing of differentiation in biasing somatosensory precursors fate toward either mechano/proprioceptive or thermo/nociceptive lineages.

## Discussion

In this study, we have addressed the not formally tested issue of the functional equivalence of the two paralogous proneural bHLH TFs Neurog1 and Neurog2, by focusing on the developing DRG. This question was not trivial, not only because of their sequence dissimilarities (Hand et al., 2005) and the phenotypic complementarities of their respective mutants (Ma et al., 1999, Venteo et al., 2019), but also because the proneural TF Ascl1 was shown unable to substitute for Neurog2 in this structure (Bertrand et al., 2002; Parras et al., 2002, Baker and Brown, 2018), likely illustrating their divergent, cell-dependent, modes of target genes regulation (Aydin et al., 2019; Vainorius et al., 2023). In contrast, we report here that replacing *Neurog2* by *Neurog1*, and vice versa, efficiently rescues the defects respectively triggered by the loss of *Neurog2* or *Neurog1* function, and that four copies of either gene are equally capable of ensuring progenitors integrity and the genesis of accurate numbers and identities of all somatosensory neuron subtypes. This establishes that *Neurog1* and *Neurog2* are functionally interchangeable in the forming DRG, implying that their respective and complementary spatio-temporal expression profiles and regulatory interactions primarily account for their apparent functional specificities, more than major differences of their biochemical properties, although subtle divergences could be uncovered in particular genetic contexts.

This is particularly well illustrated by the respective implications and efficiencies of Neurog2 and Neurog1 in suppressing the melanocyte identity. Previous findings on *Neurog1/2* single- KO and dKO have suggested a specific role for *Neurog2* in this process. They have also uncovered a developmental plasticity period -initially set between E9 and E11.5, covering the period of *Neurog2* expression- during which a group of somatosensory progenitors retains the competence to become melanoblasts (Venteo et al., 2019). Our in vivo analyses on *Neurog2^1/1^* KI and *Neurog2^1/1^;Neurog1^-/-^* KI;KO models, as well as in vitro on B16/F10 melanoblastoma cells, establish that Neurog1 does also have the ability to block the melanocyte fate. Nevertheless, the exacerbated fate switch phenotype observed in *Neurog2^1/-^* hemizygous compared to *Neurog2^+/-^* heterozygous embryos, as well as the higher doses of *Neurog1* necessary to alter the molecular identity of B16F10 cells, also indicate that *Neurog1* is less efficient than *Neurog2* in controlling this process. This appears to mainly reflects sequence dissimilarities in their bHLH domain that may influence their respective affinity for common target sequences, despite conservation of the key residues involved in DNA binding (Bertrand et al., 2002). Yet, a role for other domains outside the bHLH is also probable since the “Neurog2-bHLHNeurog1” chimeric protein exhibits a better ability than the native Neurog1 protein to down-regulate *Mitf* in vitro. Interestingly, phosphorylable amino acids in the COOH-terminal region of Neurog2, absent from Neurog1, are important for some aspects of its function (Ma et al., 2008; Hand et al., 2005). Whether these residues also influence Neurog2 efficacy to repress the melanocyte fate remains open. In addition, the striking sensory-to-melanocyte fate conversion of the *2^1neo/1neo^* transgenic model, in which exogenous Neurog1 is properly induced at E9 but prematurely down-regulated around E10.5, restricts the developmental plasticity period at E10-E11, and further supports that this phenotype mainly concerns late-migrating somatosensory progenitors, before they settle in the DRG primordium (Venteo et al., 2019; this study). This provides a rationale to the observation that in all relevant models, supernumerary *Mitf*+ melanoblasts are systematically found in the dorso- lateral migratory path of the NCC, never in the DRG anlage, and are always detected after E10.5 (Soldatov et al., 2019; Venteo et al., 2019; this study), concomitantly to the emergence of the endogenous melanoblast population (Hari et al., 2012). This likely also explains why not all somatosensory progenitors switch fate, although heterogeneity in the developmental competence among this cell population cannot be excluded. These data also provide insights into the phenotypic differences of *Neurog1^-/-^* and *Neurog2^-/-^* single-KO in this regard: 1) In *Neurog1^-/-^* KO, *Neurog2* is normally expressed in migrating somatosensory progenitors during the plasticity period, and can thus block the melanocyte fate; 2) On the opposite, in *Neurog2^-/-^*KO, Neurog1 cannot compensate for the loss of *Neurog2*, primarily because its induction in post-migratory progenitors, is incompatible with such a role, and anyway also because its onset transiently depends on Neurog2 and only occurs after E10.5 in the mutants, after the plasticity period (Ma et al., 1999; Venteo et al., 2019). Therefore, although Neurog1 has the potential to block the melanocyte fate, regulation of its spatio-temporal expression profile renders it dispensable in this process in vivo.

This study also provides definitive evidence that *Neurog1* and *Neurog2* do not differentially instruct the identity of distinct classes of somatosensory neurons in the forming DRG. This issue has been remaining ambiguous and led to heterogeneous statements within the fields of developmental and stem cell biology. The specific agenesis of most thermo/nociceptors in *Neurog1^-/-^* single-KO and the additional lack of all mechano/proprioceptors in *Neurog1^-/-^;Neurog2^-/-^*dKO, combined to their dynamic expression profiles, have suggested that *Neurog1* and *Neurog2* may participate in the specification of different fates (Ma et al., 1999; Marmigere and Ernfors, 2007; Lallemend and Ernfors, 2012; Ohayon et al., 2015). Yet, presence of few thermo/nociceptors in *Neurog1^-/-^* single-KO and of substantial numbers of mechano/proprioceptors in *Neurog2^-/-^* single-KO (Ma et al., 1999; Bachy et al., 2011; Ohayon et al., 2015; Venteo et al., 2019), as well as recent findings showing that most, if not all, NCC-derived somatosensory progenitors transiently express both TFs at some point (Venteo et al., 2019; Faure et al., 2020; Sharma et al., 2020), already challenged this view. In addition, *in vitro* strategies based on *Neurog1/2* overexpression to differentiate iPS or ES cells into distinct somatosensory neuron subtypes have yielded varied outcomes (Labau et al., 2022). In some studies, overexpression of *Neurog1* or *Neurog2* have been successfully used to specifically generate nociceptors or mechanoceptors, respectively, favoring their divergent implications (Schrenk-Siemens et al., 2014; Boisvert et al., 2015; Nickolls et al., 2020). In contrast, other studies have provided evidence that either gene can trigger the differentiation of all neuronal classes, with *Neurog2* being notably efficient in promoting the formation of thermo/nociceptors (Blanchard et al., 2015; Hulme et al., 2020; Plumbly et al., 2023). However, the use of various cell lines and (re)programming protocols (Labau et al., 2022) made it difficult to draw definitive conclusions. *In vivo* gene replacement strategies enabling functional comparisons within similar cellular environments and physiological conditions, appeared therefore particularly well suited to solve this issue.

The functional interchangeability of *Neurog1* and *Neurog2* in fate specification, along with the phenotypes of complementary transgenic mouse models in which definite temporal shifts in neurogenesis -either delayed onsets or premature arrests- have opposite effects on their DRG content, highlights the importance of differentiation timing in influencing neuronal fates: early differentiation primarily produces mechano/proprioceptors, while late differentiation mainly forms thermo/nociceptors (Figure 7). This aligns with our, and other previous birth-dating analyses establishing successive peaks of birth for both neuronal classes: E10.5-E11.5 for mechano/proprioceptors and E11.5-E12.5 for thermo/nociceptors (this study; Lawson and Biscoe, 1979; Kitao et al., 1996). However, it contrasts with another birth-dating study suggesting that the peak of genesis of thermo/nociceptors spans E10.5-E12.5, encompassing that of mechano/proprioceptors (Landy et al., 2021). Although we also found here that both neuronal classes are produced concomitantly and continuously throughout DRG neurogenesis, we observed two clear separate phases during which their respective proportions are reversed: about 85% of mechano/proprioceptors are born before E11.5, while 90% of thermo/nociceptors are born after E11.5. The reasons for these discrepancies are unclear, but the deficit of 95% of thermo/nociceptors in *Neurog1^-/-^* mutants, where neurogenesis is impaired from E11.5 onwards (Ma et al., 1999; Ohayon et al., 2015; this study), rather supports that the majority of these neurons are generated after E11.5.

Time-dependent specification and diversification of neuronal subtypes in the developing nervous system is a recurring theme, particularly well-illustrated in the retina, neocortex and ventral hindbrain. In these structures, temporal patterning mechanisms regulate the developmental potential of neural precursors over time through the sequential action of specific combinations of transcriptional regulators and environmental cues (Cepko et al., 2014; Llorca and Marin 2021; Pattyn et al., 2003; Dias et al., 2014). Functional and time- course transcriptomic analyses in the forming DRG have established that subtype diversification and maturation within both, the mechano/proprioceptive and the thermo/nociceptive lineages involve similar processes (Levanon et al., 2002; Kramer et al., 2006; Faure et al., 2020; Sharma et al., 2020; de Nooj, 2022; Chen et al., 2006; Abdo et al., 2011; Scott et al., 2011). However, the question of when these two lineages segregate remains debated. Faure et al. (2020) describe newly-born somatosensory neurons as being early committed to a particular fate. In contrast, Sharma et al. (2020) describe them as unspecified until relatively late stages, based on their initial co-expression of a set of TFs that are later maintained in specific subtypes where they control maturation features. Our functional and birth dating analyses support that the choice for a somatosensory precursor to adopt either fate is rapidly biased but depends on the developmental stage. This is consistent with the fact that populations of proprioceptive and mechanoceptive precursors, including the MafA+/cMaf+/Ret+ rapidly-adapting low-threshold mechanoreceptors, are already largely established by E11.5-E12, well before the end of DRG neurogenesis (Levanon et al., 2002; Kramer et al., 2006; Bourane et al., 2009; Luo et al., 2009; Wende et al., 2012). In the same line, Prdm12, a major determinant of the thermo/nociceptive lineage, is early detected, first in few precursor cells at E9.5-E10, and then in larger populations (Bartesaghi et al., 2019; Desiderio et al., 2019), paralleling the gradual increase in the production of this neuronal class over time. The mechanisms underlying this time-dependent lineage segregation remain however unclear. Evidence indicates that somatosensory progenitors as well as young post- mitotic precursors, are not intrinsically and irreversibly fated but exhibit developmental plasticity. Indeed, in a mouse model where late-migrating NCC-derived somatosensory progenitors are specifically depleted, early progenitors mainly generate mechano/proprioceptors when they normally differentiate at E9.5-E11, but mostly produce thermo/nociceptors when they atypically differentiate at E11-E13.5 (Ohayon et al., 2015; Venteo et al., 2019), further underscoring the importance of the differentiation timing. Moreover, forced expression of the neurotrophin receptor TrkC from the *TrkA* locus, can induce some TrkA+ nociceptive precursors to become proprioceptors (Moqrich et al., 2004). Together, these data suggest key roles for extrinsic instructive signals emanating from the rapidly evolving environment around the DRG anlagen (de Nooj, 2022). Yet, the continuous production of both neuronal classes throughout neurogenesis (Landy et al., 2021; Kitao et al., 1996; this study) also suggests that these signals might be active concomitantly, with their respective intensities and/or the capacity of precursor cells to respond varying over time. This simultaneous activity could also explain the temporary co-expression of several genes encoding subtype-restricted maturation TFs (Sharma et al., 2020). Alternatively, this may reflect early active pan-somatosensory regulatory elements in their promoter regions, later overridden by subtype-specific elements, ultimately limiting their expression at stages when their function is engaged.

As proneural TFs, Neurog1 and Neurog2 also influence cell proliferation by notably promoting cell cycle exit and activating the Notch signaling pathway which is critical for maintaining the integrity of undifferentiated proliferative precursors throughout neurogenesis (Bertrand et al., 2002; Hu et al., 2011). Their respective expression, which roughly coincides with the two successive neurogenic phases of the DRG generating, respectively, about 25% and 75% of the somatosensory neurons, raised the possibility that they may actively and differently control the distinct proliferative properties of early- and late-migrating progenitors (Lawson and Biscoe, 1979; Franck and Sanes, 1991; Ma et al., 1999; Marmigere et al., 2007; Ohayon et al., 2015; this study). Nevertheless, the normal numbers of neurons in the DRG of *Neurog1^2/2^*and *Neurog2^1/1^* single-KI models establish that both TFs are also essentially interchangeable in this process and, therefore, do not account for the distinct contributions of the different progenitor pools. This result, together with the observation that early progenitors consistently form about 20% of the DRG neurons regardless of whether they differentiate between E9.5-E11 or E11- E13.5 (Ohayon et al., 2015; Venteo et al., 2019), suggests that their proliferative capacities are early intrinsically fixed, in contrast to their fate.

Functional equivalence of *Neurog1* and *Neurog2* in the developing DRG raises the issue of the evolutionary rationale for having selected and maintained two paralogous genes with similar functions. It has been proposed that duplication of proneural genes followed by subsequent modifications of their coding sequences and/or regulatory elements, have contributed to the complexification and diversification of the nervous system (Furlong and Graham, 2005; Baker and Brown, 2018). In this regards, it is interesting to note that the development of the zebrafish DRG which contain only few neurons, is controlled by a single *Neurogenin* family member orthologous to *Neurog1* (Raible and Eisen 1992; MacGraw et al., 2008). It is thus conceivable that in higher Vertebrates, after duplication of an ancestral gene, the regulatory elements of *Neurog1* and *Neurog2* have evolved independently of each other, and along with the diversification and expansion of the populations of NC-derived progenitors, allowing the formation of increased numbers of neurons and the emergence of new functional identities. In this context, our findings suggest a model in which, regardless of their nature, combined expression of *Neurog1* and *Neurog2* allows a continuum of neurogenesis between E9.5 and E13.5, so that early –low-proliferating–, and late – highly- proliferating– progenitor populations differentiate during successive time-windows, and within distinct environments, that determine their fate as either mechano/proprioceptors or thermo/nociceptors, thus underlying the relative proportions of 20% and 80% of both neuronal classes. In this model, shortening the periods of *Neurog1/2* expression in a way or another, consequently restricts neurogenesis to defined periods with predictable impacts on DRG content.

Beyond the spinal peripheral somatosensory system, *Neurog1* and *Neurog2* also control neurogenesis in several other regions of the nervous system, either individually, like in the ventral midbrain dopaminergic nuclei (Anderson et al., 2006; Kele et al., 2006), or on a combinatorial mode, like in the olfactory system (Shaker et al., 2012; Dixit et al., 2014) and cranial sensory ganglia (Ma et al., 1998; Fode et al., 1998). Preliminary evidences suggest that *Neurog1* can efficiently replace *Neurog2* during the genesis of midbrain dopaminergic neurons (data not shown). Similarly, the complementary defects observed in the cranial sensory ganglia of *Neurog1^-/-^* and *Neurog2^-/-^*single-KO are also rescued in the single-KI models (data not shown). It is noteworthy that the latter result was expected since in the chick embryo, the spatio-temporal dynamic of *Neurog1* and *Neurog2* expression in cranial sensory progenitors originating from the neurogenic placodes and the neural crest, are strikingly inverted compared to the mouse (Begbie et al., 2002; Furlong and Graham, 2005). It will be interesting to evaluate other *Neurog1/2*-dependent neural structures to determine whether their functional specificities generally rely solely on their complementary expression profiles.

## Acknowledgments

We thank F. Guillemot for Neurog mutants, Q. Ma, JF. Brunet and E. Bellefroid for materials, and M. Simonelig and A. Chartier for technical help. This work was supported by INSERM, France; CNRS, France; Fondation pour la Recherche Medicale, France; Agence Nationale pour la Recherche, France; and University of Montpellier, France.

## Author contributions

S.D., P.Cabochette, S.V. and A.P. performed the experiments. S.D., P.Cabochette., A.P., P.Carroll and F.A. designed the experiments and interpreted the data. S.D., P.Cabochette and A.P. wrote the manuscript.

