## Supplementary Figures and Legends for "Functional Equivalence of the Proneural Genes *Neurog1* and *Neurog2* in the Developing Dorsal Root Ganglia Highlights the Importance of Timing of Neurogenesis in Biasing Somatosensory Precursor Fates"

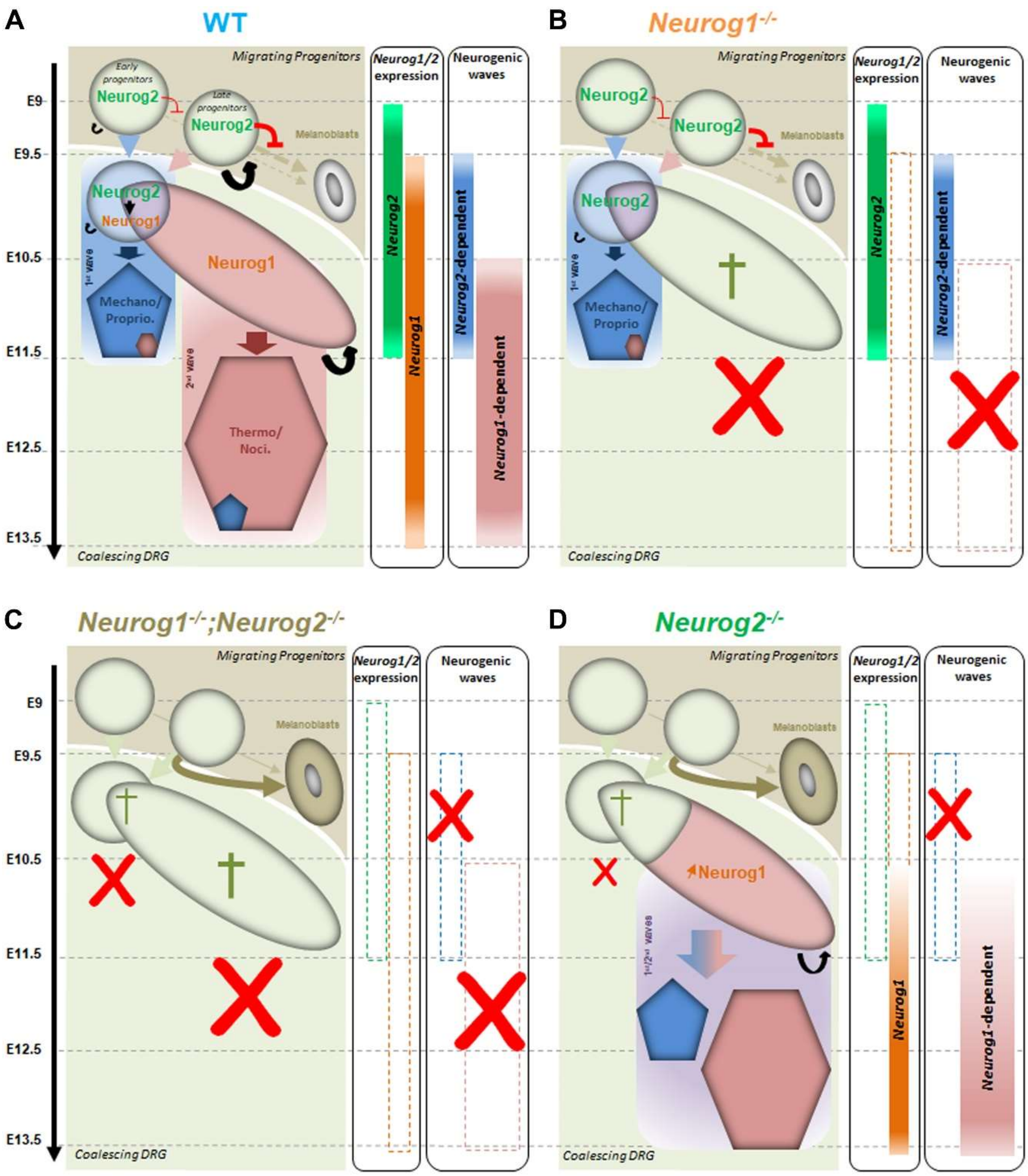


**Supplementary Figure 1. Schematic illustration of the dynamic expression profiles and combinatorial roles of *Neurog1* and *Neurog2* during DRG neurogenesis**

**A.** In WT, *Neurog2* and *Neurog1* exhibit dynamic and complementary expression profiles which correlate with the two main successive waves of neurogenesis underlying DRG development. These waves are classically assumed to mainly generate mechano/proprioceptors (Mechano/proprio.) and thermo/nociceptors (Thermo/noci.), respectively, from early migrating –low proliferating (thin curved arrows)– and late migrating –highly proliferating (thick curved arrows)– progenitor populations. Note that *Neurog2* is induced in all somatosensory progenitors at some point where it notably represses the melanocyte fate and transiently controls onset of *Neurog1* expression in post-migratory progenitors. See text for details.

**B.** In *Neurog1*$^{-/-}$ KO, the second neurogenic wave is specifically abolished (red crosses), resulting in agenesis of most thermo/nociceptors and apoptosis of the late progenitor population (green cross). In contrast, the first neurogenic wave proceeds normally, in line with the normal expression of *Neurog2* from E9 to E11.5.

**C.** In *Neurog1*$^{-/-}$;*Neurog2*$^{-/-}$ dKO, both neurogenic waves are impaired (red crosses) and no somatosensory neuron is ever produced. In these mutants, somatosensory progenitors either die through apoptosis (green crosses) or switch fate to become melanoblasts (brown arrows).

**D.** In *Neurog2*$^{-/-}$ KO, onset of neurogenesis (red crosses) and of *Neurog1* expression are delayed by 24h. During this transient period, pools of somatosensory progenitors either die (green cross) or become melanoblasts (brown arrows). From E10.5, however, *Neurog1* is eventually induced and neurogenesis is initiated, albeit from a reduced reservoir of progenitors. This leads to a reduction of all neuronal classes. See text for details.

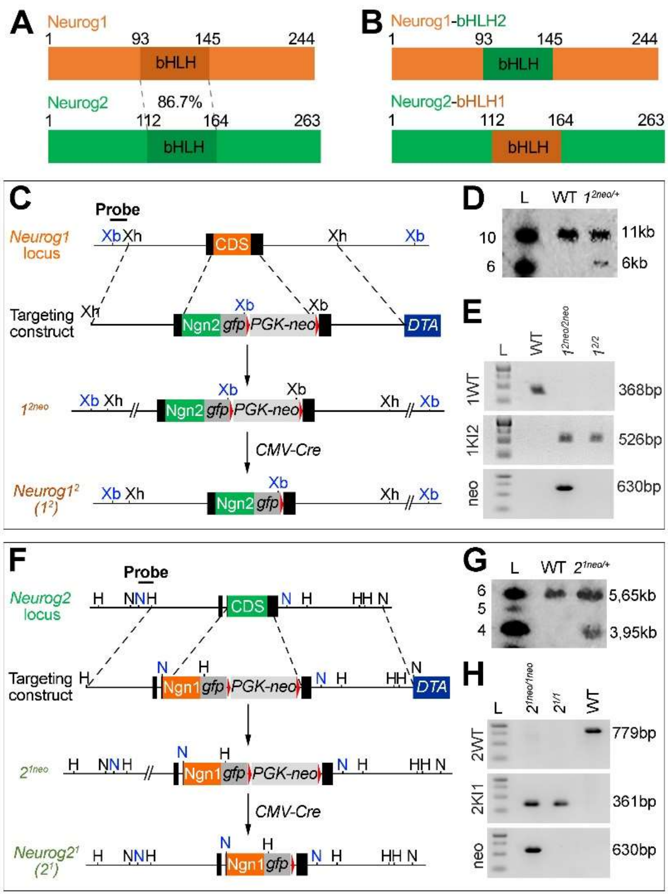


**Supplementary Figure 2. Generation of *Neurog1*$^{KINeurog2}$ and *Neurog2*$^{KINeurog1}$ Knock-in alleles**

**A.** Schematic representation of the Neurog1 and Neurog2 proteins illustrating their respective sizes and the 86.7% of identity within their bHLH domains.

**B.** Schematic representation of chimeric proteins in which the bHLH domain of Neurog2 has been inserted into the Neurog1 protein (Neurog1-bHLH2) and conversely (Neurog2-bHLH1).

**C.** Schematic illustration of the strategy used to generate the *Neurog1*$^{2neo}$ ($1^{2neo}$) and *Neurog1*$^{2}$ alleles. The *Neurog1* locus contains one single exon harboring the entire coding sequence (CDS). The targeting construct consisted in: (i) a 5'-arm of homology, between an upstream XhoI site and the ATG codon; (ii) a sequence containing the Neurog2 CDS followed by an ires-GFP and a Floxed-PGK-neo cassette; and (iii) a 3'-arm of homology, between the Stop codon and a downstream XhoI site. After electroporation, selection and screening, recombined ES cells in which the CDS of Neurog1 was replaced by the "Neurog2CDS-iresGFP-PGKneo" cassette (see D) were used to first establish the *Neurog1*$^{2neo}$ (*1*$^{2neo}$) mouse line. This line was then crossed with a CMV-Cre line to remove the Floxed-PGK-neo cassette and establish the *Neurog1*$^{2}$ which represent a "clean" Knock-in allele of the gene.

**D.** Illustrative image of a Southern Blot analysis illustrating homologous recombination in ES cells. DNAs from electroporated ES clones was extracted, digested with XbaI, migrated on an agarose gel and transferred onto a Nylon membrane that was hybridized with a $P^{32}$-labelled DNA probe (XbaI-XhoI fragment) located upstream of the targeted region. The probe revealed a 11kb fragment in the WT allele and a 6kb fragment in the recombined allele due to the presence of an exogenous XbaI site in the cassette.

**E.** Illustrative images of PCR diagnostics of WT, *Neurog1*$^{2neo/2neo}$ (*1*$^{2neo/2neo}$) and *Neurog1*$^{2/2}$ (*1*$^{2/2}$) animals. Both transgenic homozygous animals lack the 368bp WT fragment but contained the 526bp KI fragment. *Neurog1*$^{2/2}$ animals additionally lack the 630bp neo fragment amplified in *Neurog1*$^{2neo/2neo}$.

**F.** Schematic illustration of the strategy used to generate the *Neurog2*$^{1neo}$ (*2*$^{1neo}$) and *Neurog2*$^{1}$ alleles. The *Neurog2* locus contains two exons but only is coding. The targeting construct consisted in: (i) a 5'-arm of homology, between an upstream HindIII site and the ATG codon; (ii) a sequence containing the Neurog1 CDS followed by an ires-GFP and a Floxed-PGK-neo cassette; and (iii) a 3'-arm of homology, between the Stop codon and a downstream NsiI site. After electroporation, selection and screening, recombined ES cells in which the CDS of Neurog2 was replaced by the "Neurog1DS-iresGFP-PGKneo" cassette (see G) were used to first establish the *Neurog2*$^{1neo}$ (*2*$^{1neo}$) mouse line. This line was then crossed with a CMV-Cre line to remove the Floxed-PGK-neo cassette and establish the *Neurog2*$^{1}$ line

**G.** Illustrative image of a Southern Blot analysis illustrating homologous recombination in ES cells. DNAs from electroporated ES clones were extracted, digested with NsiI, migrated on an agarose gel and transferred onto a Nylon membrane that was then hybridized with a $P^{32}$-labelled

DNA probe (NsiI-HindIII fragment) located upstream of the targeted region (see F). The probe revealed a 5.65kb fragment in the WT locus and a 3.95kb fragment in the recombined locus due to the presence of an exogenous NsiI site in the cassette.

**H.** Illustrative images of PCR diagnostics of WT, *Neurog2*$^{1neo/1neo}$ (*2*$^{1neo/1neo}$) and *Neurog2*$^{1/1}$ (*2*$^{1/1}$) animals. Both transgenic homozygous animals lack the 779bp WT fragment but contained the 361bp KI fragment. *Neurog2*$^{1/1}$ animals additionally lack the 630bp neo fragment amplified in *Neurog1*$^{2neo/2neo}$.

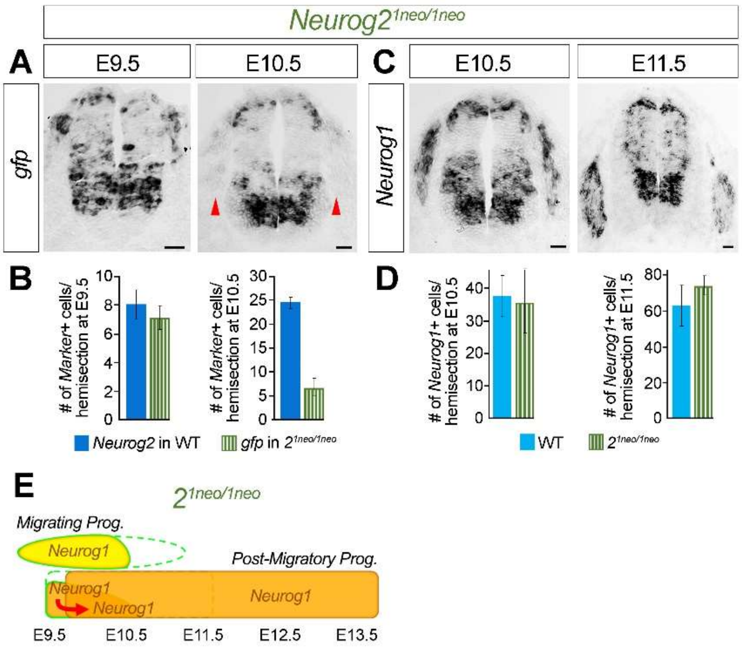


**Supplementary Figure 3. Presence of the PGK-neo cassette in *Neurog2*$^{1neo/1neo}$ animals alters the activity of the endiogenous locus *Neurog2* locus.**

**A.** Representative images of *in situ* hybridization for *gfp* at E9.5 and E10. on transverse thoracic spinal cord sections of *Neuro2*$^{1neo/1neo}$ embryos. Red arrowheads point to the coalescing DRG that abnormally express low levels of *Neurog1* in E10.5 *Neurog2*$^{1neo/1neo}$ embryos. Scale bar, 50µm.

**B.** Quantitative analysis of the number of *Neurog2*+ cells in WT (dark blue columns) and *gfp*+ cells in *Neurog2*$^{1neo/1neo}$ (green striped columns) embryos at E9.5 and E10.5, as indicated, showing premature down-regulation of *gfp* in the transgenic animals.

**C.** Representative images of *in situ* hybridization for *Neurog1* at E10.5 and E11.5 (B) on transverse thoracic spinal cord sections of *Neuro2*$^{1neo/1neo}$ embryos. Scale bar, 50µm.

**D.** Comparative quantitative analysis of the number of *Neurog1*+ cells in WT (blue columns) and *Neurog2*$^{1neo/1neo}$ (green striped columns) embryos at E10.5 and E11.5, as indicated, showing no significant difference.

**E.** Schematic summary of the dynamic expression profiles of *Neurog1* (orange colors) in migrating (oval frame) and/or post-migratory (rectangle frame) somatosensory progenitors in *Neurog2*$^{1neo/1neo}$ embryos between E9.5 and E13.5. In this model, exogenous *Neurog1* expression from the *Neurog2* locus (mimicked by the reporte gene *gfp*) is initiated on time at E9 but is prematurely down-regulated at E10.5. Nevertheless, the early expression of *Neurog1* appears sufficient to induce the endogenous *Neurog1* locus, which is subsequently normally expressed over time.

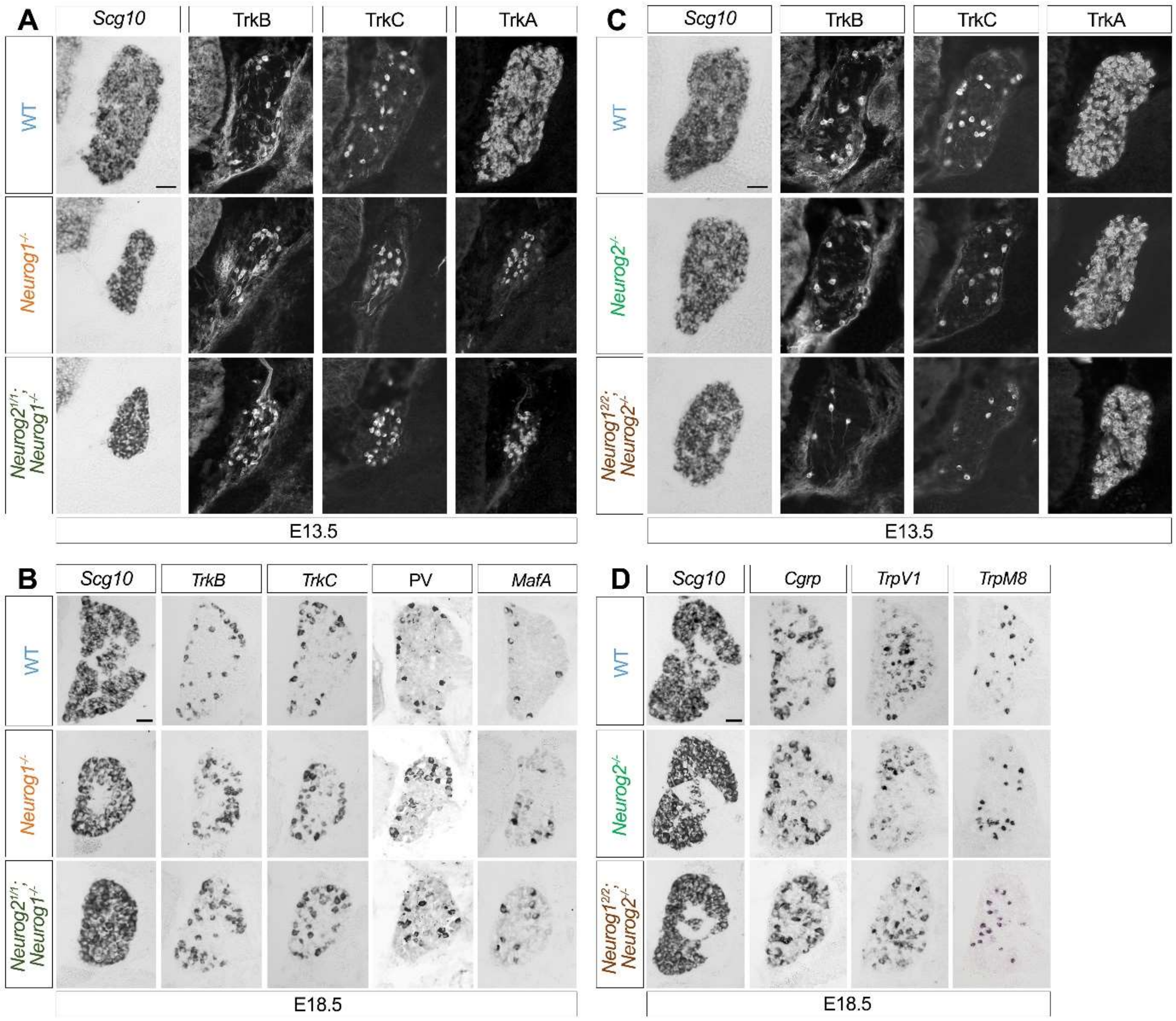


**Supplementary Figure 4. Histological analysis of *Neurog1*$^{-/-}$, *Neurog2*$^{1/1}$*;Neurog1*$^{-/-}$, *Neurog2*$^{-/-}$, *Neurog1*$^{2/2}$*;Neurog2*$^{-/-}$ embryos at E13.5 and E18.5.**

**A.** Representative images of *in situ* hybridization for *Scg10*+ or immunofluorescent staining for TrkB, TrkC and TrkA on transverse DRG sections of WT, *Neurog1*$^{-/-}$ KO and *Neurog2*$^{1/1}$;*Neurog1*$^{-/-}$ KI;KO embryos at E13.5. Scale bar, 50µm.

**B.** Representative images of *in situ* hybridization for *Scg10*+, *TrkB*, *TrkC* and *MafA* or immunofluorescent staining for PV on transverse DRG sections of WT, *Neurog1*$^{-/-}$ KO and *Neurog2*$^{1/1}$;*Neurog1*$^{-/-}$ KI;KO embryos at E18.5. Scale bar, 50µm.

**C.** Representative images of *in situ* hybridization for *Scg10* or immunofluorescent staining for TrkB, TrkC and TrkA on transverse DRG sections of WT, *Neurog2*$^{-/-}$ KO and *Neurog1*$^{2/2}$;*Neurog2*$^{-/-}$ KI;KO embryos at E13.5. Scale bar, 50µm.

**D.** Representative images of *in situ* hybridization for *Scg10*, *Cgrp*, *TrpV1* and *TrpM8* on transverse DRG sections of WT, *Neurog2*$^{-/-}$ KO and *Neurog1*$^{2/2}$;*Neurog2*$^{-/-}$ KI;KO embryos at E18.5. Scale bar, 50µm.

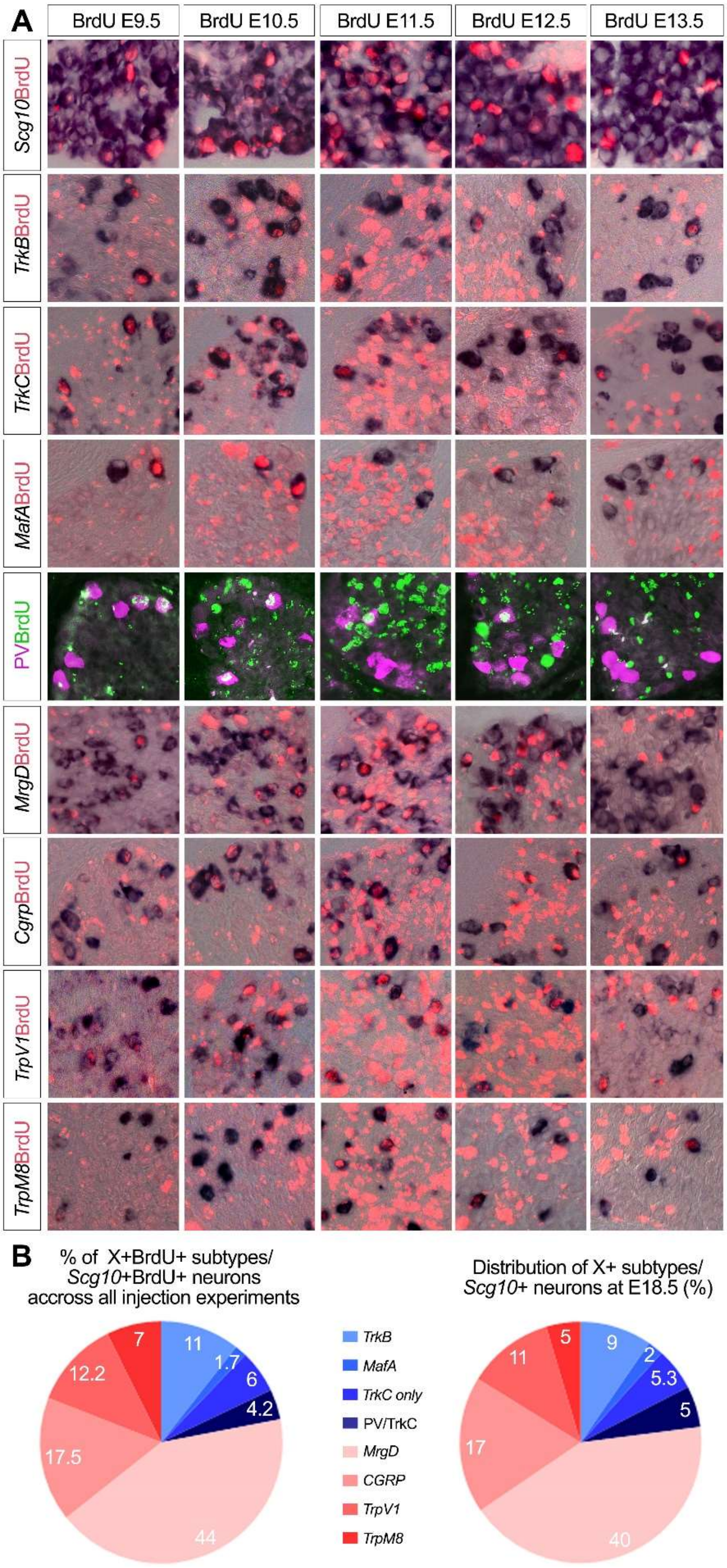


**Supplementary Figure 5. Birth-dating analysis of mechano/proprioceptive and thermo/nociceptive subtypes.**

**A.** Representative images of combined immunostaining for BrdU and *in situ* hybridization for *Scg10*, *TrkB*, *TrkC*, *MafA*, *MrgprD*, *Cgrp, TrpV1* or *TrmP8*, and double-immunofluorescent staining for BrdU and PV on transverse DRG sections of E18.5 WT embryos injected with BrdU at the indicated stages.

**B.** Pie Charts showing either the proportions of the total numbers of each X+/BrdU+ mechano/proprioceptive (blue colors) and thermo/nociceptive (reddish colors) subtypes relative to the total number of *Scg10*+BrdU+ accross all experiments (left panel) or the proportions of each X+ subtypes relative to the total number of *Scg10*+ DRG neurons counted in E18.5 embryos (right panel). No major difference was observed between the two quantifications, indicating that all X+ subtypes were properly represented in our experimental paradigm.
